# Analyzing kinetic signaling data for G-protein-coupled receptors

**DOI:** 10.1101/2020.01.20.913319

**Authors:** Sam R.J. Hoare, Paul H. Tewson, Anne Marie Quinn, Thomas E. Hughes, Lloyd J. Bridge

## Abstract

In classical pharmacology, bioassay data are fit to general equations (e.g. the dose response equation) to determine empirical drug parameters (e.g. EC_50_ and E_max_), which are then used to calculate chemical parameters such as affinity and efficacy. Here we used a similar approach for kinetic, time course signaling data, to allow empirical and chemical definition of signaling by G-protein-coupled receptors in kinetic terms. Experimental data are analyzed using general time course equations (model-free approach) and mechanistic model equations (mechanistic approach) in the commonly-used curve-fitting program, GraphPad Prism. A literature survey indicated signaling time course data usually conform to one of four curve shapes: the straight line, association exponential curve, rise-and-fall to zero curve, and rise-and-fall to steady-state curve. In the model-free approach, the initial rate of signaling is quantified and this is done by curve-fitting to the whole time course, avoiding the need to select the linear part of the curve. It is shown that the four shapes are consistent with a mechanistic model of signaling, based on enzyme kinetics, with the shape defined by the regulation of signaling mechanisms (e.g. receptor desensitization, signal degradation). Signaling efficacy is the initial rate of signaling by agonist-occupied receptor (*k*_*τ*_), simply the rate of signal generation before it becomes affected by regulation mechanisms, measurable using the model-free analysis. Regulation of signaling parameters such as the receptor desensitization rate can be estimated if the mechanism is known. This study extends the empirical and mechanistic approach used in classical pharmacology to kinetic signaling data, facilitating optimization of new therapeutics in kinetic terms.

## Introduction

Pharmacological data analysis provides the drug activity parameters used to optimize the biological activity of new therapeutics and to identify mechanisms of receptor-mediated signal transduction for G-protein-coupled receptors (GPCRs). In the typical analysis process, activity is measured at various drug concentrations and the data fit to a concentration-response equation by curve fitting, for example to a sigmoid curve equation ^1-3^. This yields empirical drug parameters, usually the potency (EC_50_) and a measure of the maximal signaling capacity (E_max_). This analysis is described as “Model-free” because it makes minimal assumptions regarding the mechanism of signal transduction. These parameters can then be used to calculate mechanistic drug parameters that define the effect of the drug on the system in chemical terms (e.g. efficacy and macroscopic affinity) ^3-8^. These chemical parameters aid the translation of drug effect measurement from in vitro test systems to animal models and human disease, for example by minimizing the effect of tissue-specific parameters on the quantification of drug activity. Biased agonism measurement is an example of this process currently in use by many laboratories. Signaling assay data are fit to concentration-response equations and the empirical parameters EC_50_ and E_max_ used to calculate mechanistic parameters of biased signaling, such as transducer coefficients and log efficacy ratios (reviewed in ^9^).

The manner in which GPCR signaling changes over time, i.e. the kinetics / dynamics of response, is currently of considerable interest in quantitative pharmacology. The time course of GPCR response has been measured since the earliest studies of muscle contraction ^1^, through the discovery of G-protein-mediated signaling ^10^, the application of Ca^2+^ indicator dyes ^11^, and the most recent advances in biosensor technology that enable high-throughput signaling kinetic analysis ^12-14^. Measuring the time course of receptor signaling revealed fundamental mechanisms of GPCR function. The classic rise-and-fall time course of cAMP in S49 cells indicated desensitization of β_2_ adrenoceptor signaling ^15^. Sustained signaling after agonist washout by parathyroid hormone, sphingosine 1-phosphate and thyroid-stimulating hormone receptors revealed signaling by internalized receptors ^16-18^, a new spatiotemporal paradigm of GPCR signaling of potential therapeutic utility ^19-23^. Signaling kinetics can also affect quantification of drug activity. Of potential concern, drug effect can be dependent on the time point at which the response is measured. For example, biased signaling of the D_2_ dopamine receptor was shown to be time-dependent ^24^. For the CB_1_ receptor, the potency for internalization increased over time ^25^. For AT_1_ angiotensin receptor-mediated arrestin recruitment the potency increased over time, as did the E_max_ of partial agonists ^26^.

A data analysis framework for time course data of GPCR signaling would enable the quantification of efficacy in kinetic terms. One way to do this is to measure the initial rate of signaling, analogous to the initial rate of enzyme activity ^26-31^, an approach recently applied to receptor receptor-arrestin interaction ^26^. Kinetic analysis could also enable measurement of regulation of signaling parameters, such as the rate of receptor desensitization. At present, time course data for GPCR signaling are rarely analyzed by curve fitting to estimate kinetic pharmacological parameters (in contrast to the near-universal application of curve fitting to concentration-dependence data). The absence of curve fitting to time course data might be resulting in lost opportunities, particularly for biosensor assay modalities in which the time course data are collected by default. Kinetic insights into GPCR function might be being missed. Potential benefits of the kinetic dimension to ligand optimization could be going unrealized. The goal of this study was to develop a straightforward data analysis framework for quantifying the kinetics of GPCR signaling, enabling investigators to measure kinetic drug parameters useful for practical pharmacological application.

## Results

In this study, a data analysis framework for curve fitting of time course data was developed for GPCR signaling kinetics/dynamics. First, we identified the types of time course curve shape by conducting a comprehensive literature survey. It was discovered that the curve shapes, more precisely the temporal profiles, conform to a limited number of shapes (four). This survey also showed how the time course is shaped by regulation of signaling mechanisms (e.g. receptor desensitization and response degradation). The shapes were defined by simple equations that can be used to analyze time course data using familiar curve-fitting software (e.g. Prism, GraphPad Software, Inc., San Diego CA). This analysis yields a kinetic drug parameter, the initial rate of signaling by agonist-occupied receptor, that defines efficacy as the rate of signal generation before it is impacted by regulation of signaling mechanisms. This parameter, termed *k*_*τ*_, provides a biologically meaningful and intuitive metric of the kinetics of signal transduction. Finally, mechanistic models are applied to the time course data using a kinetic mechanistic model of agonism developed previously ^32^, and extended here to incorporate receptor desensitization and sustained signaling by internalized receptors. This analysis demonstrated the model-free equations emerge as general forms of the mechanistic equations, providing a mechanistic foundation for the analysis approach.

### Literature survey of time course shapes

The first step in analyzing biological activity data is visual inspection of the data. In pharmacological data analysis, this led to the realization that concentration-response data are usually described by a sigmoid curve (when the x axis is the logarithm of ligand concentration) ^1,2^. Here we surveyed the GPCR signaling literature for time course data. The survey was designed to be comprehensive, spanning 1) The full range of heterotrimeric G-protein classes (Gs, Gi, Gq/11, G12/13); 2) response times of milliseconds to hours; 3) Proximity to the receptor, from direct receptor interaction (G-protein and arrestin) to downstream signals (e.g. cell contraction/relaxation and gene expression); 4) Types of response, including chemical (second messengers), mechanical (cell structure change), electrical (ion channel currents) and genetic (gene expression). This was done for experiments when the receptor was exposed continuously to the agonist, i.e. agonist washout experiments were not considered.

Within this survey, the large majority of time course profiles conformed to one of four shapes (Fig. 1). Each shape is defined by a corresponding equation (equations 1 to 4 in Methods). The equations are amenable to straightforward curve-fitting analysis in familiar software. The shapes are, in order of increasing complexity:

**Figure 1.**
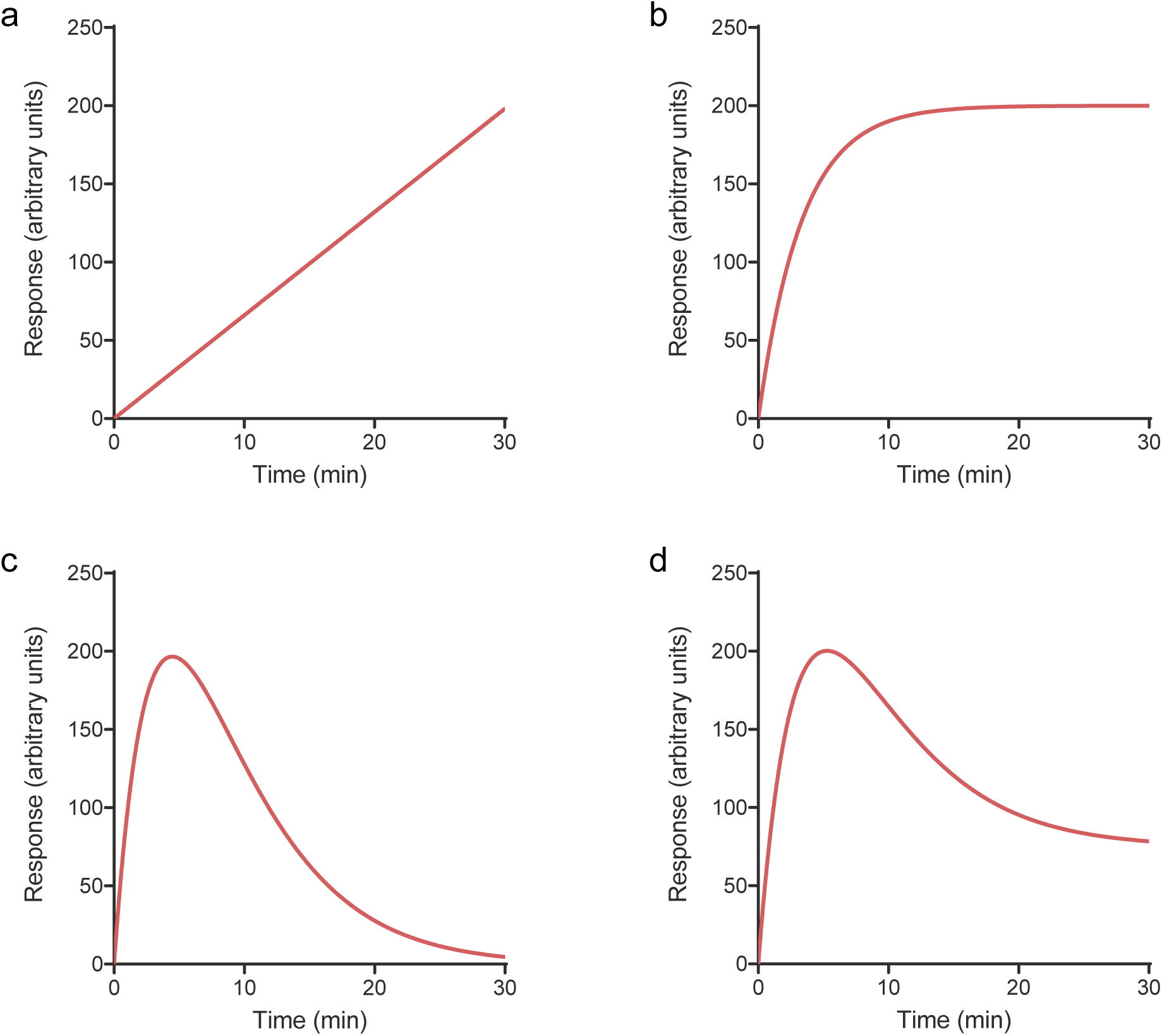
Time course profile shapes typically observed for GPCRs. (a) Straight line. (b) Association exponential curve. (c) Rise-and-fall to baseline curve. (d) Rise-and-fall to steady-state curve. Data for (a) were simulated using equation (1) (Slope, 6.6 response units.min^-1^), (b) with equation (2) (SSR, 200 response units; *k*, 0.30 min^-1^), (c) with equation (3) (*C*, 120 response units.min^-1^; *k*_1_, 0.25 min^-1^; *k*_2_, 0.20 min^-1^) and (d) with equation (4) (SSR, 75 response units; *D*, 25 (note *D* is unitless); *k*_1_, 0.25 min^-1^; *k*_2_, 0.20 min^-1^). The Baseline parameter in each equation was set to zero.

1. The straight line (Fig. 1a). Response increases continuously over time at a constant rate (defined by equation (1)).
2. The association exponential curve (Fig. 1b, equation (2)). Initially, response increases rapidly, being almost linear. The response then slows then finally approaches a plateau at which the response level remains constant over time.
3. The rise-and-fall to baseline curve (Fig. 1c, equation (3)). The first phase resembles the association exponential curve – response increases rapidly at first, then slows. The response then reaches a peak level. Subsequently, the response level decreases and finally falls back to the baseline level, i.e. the level before addition of ligand.
4. The rise-and-fall to steady-state curve (Fig. 1d, equation (4)). Response rises and falls as for the rise-and-fall to baseline curve. However, the response declines to a steady-state level, above that of the baseline level before addition of ligand.

### Straight line time course profile

The straight line profile was evident in second messenger responses under a specific condition – when signaling was unregulated, i.e. when regulation of signaling mechanisms were blocked (Fig. 2). GPCR signaling is regulated over the short term by two primary mechanisms: receptor desensitization (involving receptor phosphorylation and subsequent arrestin binding) ^33-35^ and degradation of the response (for example metabolism of second messengers) ^36-38^. The straight line profile was observed when both mechanisms were blocked. For example, AT_1_ angiotensin receptor-stimulated inositol phosphates (IP) production was linear when desensitization was blocked (using arrestin knock out cells) and response degradation was blocked (using Li^+^ to block IP breakdown) (Fig. 2a) ^39^. The same result was obtained for the proteinase-activated receptor PAR1 receptor (see Fig. 2a in ^40^). A second example is provided by the GnRH_1_ receptor, which lacks a C-terminal tail and so does not interact with arrestin ^41^. A linear time course was observed, in the presence of Li^+^ (Fig. 2b) ^42^. (See also Fig. 1a,b in ^43^ and Fig. 3 in ^44^.) A third example is provided by the glucagon receptor. A linear time course of cAMP accumulation was observed in hepatic membranes ^45^. The membranes lacked cAMP phosphodiesterase activity ^46^, and likely lacked receptor desensitization components owing to extensive washing of the preparation (Fig. 2c).

**Figure 2.**
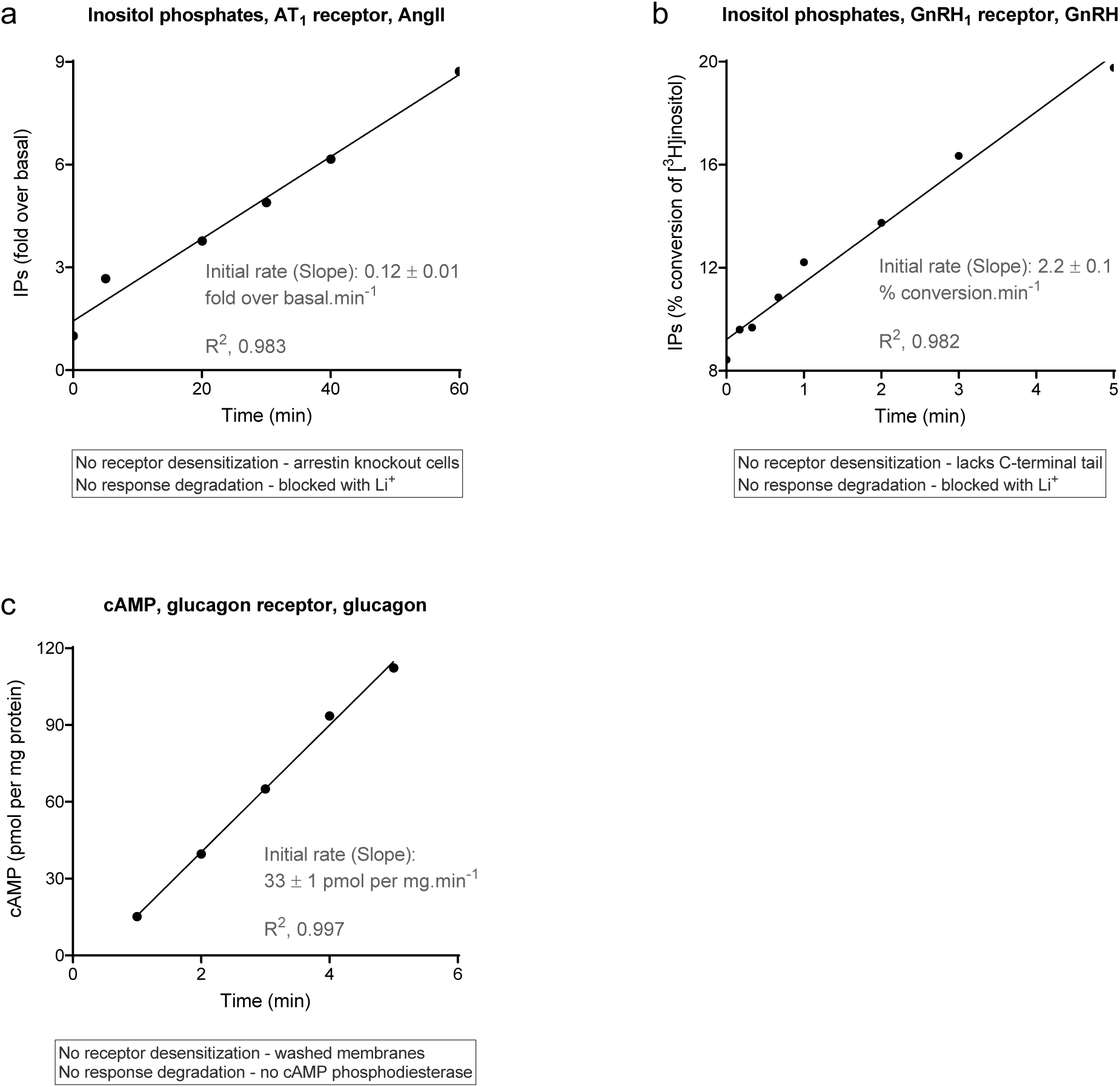
Linear time course profile examples of GPCR signaling. In all cases, regulation of signaling activity (receptor desensitization and response degradation) was minimized as described in the text boxes. (a) Inositol phosphates production via the AT1 receptor stimulated by 100 nM AngII in fibroblasts from arrestin knock-out mice (from Fig. 3c of ^35^). (b) Inositol phosphates production via the GnRH1 receptor stimulated by 1 μM GnRH in HEK293 cells (from Fig. 4a of ^38^). (c) cAMP production in hepatic membranes stimulated by 2 μM glucagon (data from Fig. 1 of ^41^, basal response in the absence of glucagon subtracted). Note the membrane preparation used lacks cAMP phosphodiesterase activity 42. Data were fit to the straight line equation (equation (1)) using Prism 8 ^89^. The Slope value on the panels, which is equal to the initial rate and *k*_*τ*_, is the fitted value ± the fit SEM ^93,96^.

**Figure 3.**
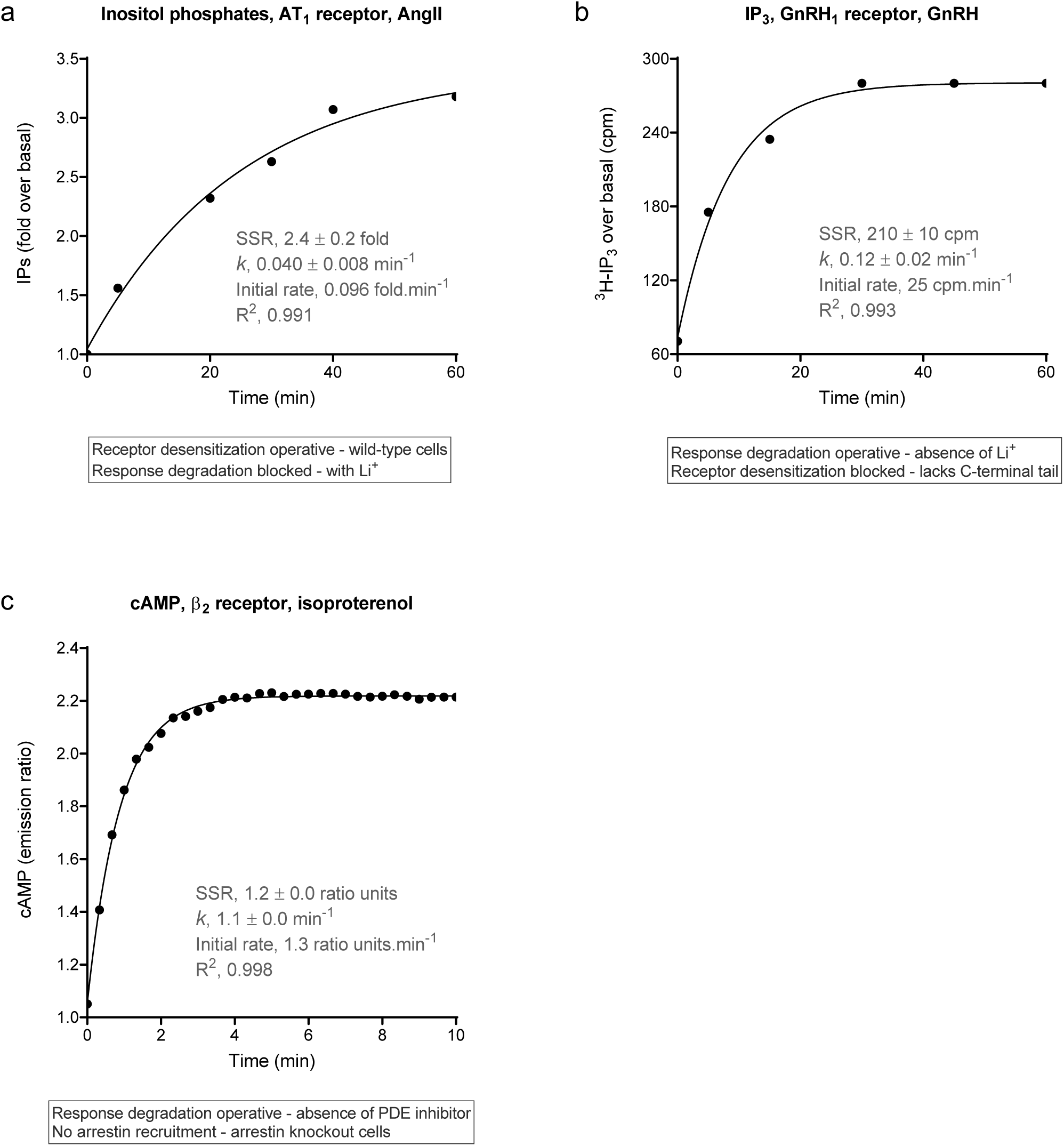
Association exponential time course profile examples of GPCR signaling. In all cases, one of the regulation of signaling mechanisms was in operation and one was blocked (see text boxes). Data were fit to the association exponential equation (equation (2)) using Prism 8 ^90^. (a) Inositol phosphates production via the AT1 receptor stimulated by 100 nM AngII in fibroblasts from wild-type mice (from Fig. 3c of ^35^). (b) Inositol phosphates production via the GnRH1 receptor stimulated by 85 nM GnRH in rat granulosa cells in the absence of Li^+^ (from Fig. 3 of ^40^). (c) cAMP production via the β2 adrenoceptor stimulated by 1 μM isoproterenol in fibroblasts from arrestin knock-out mice (data from Fig. 5d of ^45^). Note a slight decline was observed at later time points (> 10 min), which have been excluded from the figure. The fitted values SSR (steady-state response) and *k* (the rate constant) are the fitted value ± the fit SEM ^93,96^. The initial rate, equal to *k*_*τ*_, was calculated as the steady state response (SSR) multiplied by the rate constant *k*.

Linear time course data were also observed for long duration responses, far downstream, at the level of DNA, specifically gene expression and DNA synthesis (Supplementary Fig. S1) ^47,48^. In these responses there was a delay before the onset of the response. This has been rationalized for gene expression response by a need for a build-up of signal transduction intermediates to a threshold level that then initiates the process ^47^.

### Association exponential time course profile

The association exponential profile was the most commonly-observed time course shape, especially for second messenger molecules such as cAMP and inositol phosphates, but also for a variety of other signals, from upstream events such as arrestin recruitment to downstream cellular functions such as muscle cell relaxation (Fig. 3, Supplementary Fig. S2). Regarding regulation of signaling, the profile was evident when one of the two mechanisms was operative and one was blocked (the mechanisms being receptor desensitization and response degradation). Examples include AT_1_ receptor-stimulated IP production, with receptor desensitization in operation (arrestin wild-type cells) but without response degradation (IP metabolism blocked by Li^+^) (Fig. 3a) ^39^; GnRH_1_ receptor-simulated IP production with response degradation in operation (no Li^+^ present) but without receptor desensitization (the GnRH receptor lacking a C-terminal tail) (Fig. 3b) ^44^; and β2 adrenoceptor-stimulated cAMP generation with response degradation (cAMP metabolism) but without receptor desensitization (arrestin knockout cells) (Fig. 3c) ^49^.

The association exponential profile was observed in direct receptor-transducer interaction assays, including receptor-G-protein interaction and receptor-arrestin interaction (Supplementary Fig. S2a,b) ^26,50^. This was expected; mechanistically, the response is likely resultant from a bimolecular interaction and the association exponential equation is a general form of the familiar bimolecular association kinetics equation. G-protein activation was also described by the profile, measured by [^35^S]GTPγS binding, or by a G-protein activation biosensor (Supplementary Fig. S2c,d) ^50,51^. cAMP production responses also conform to the profile, both stimulation (Supplementary Fig. S2e) and inhibition (Supplementary Fig. S2f) ^52,53^. A specialized signaling pathway, β-catenin stabilization via the Wnt-Frizzled system, was also described by the profile ^54^ (Supplementary Fig. S2g). Electrical signaling was also described by the association exponential profile, as shown by GIRK channel gating in *Xenopus* oocytes (Supplementary Fig. S2h) ^55^. In the final example, a downstream cellular response was found to conform to the profile, relaxation of human airway smooth muscle cells via the β_2_ adrenoceptor (Supplementary Fig. S2i) ^56^.

### Rise-and-fall to baseline time course profile

The rise-and-fall to baseline profile is a classic curve shape in pharmacology, leading to the definition of “Fade” (decline in the response to a continuous application of agonist ^57^) and the discovery of the underlying mechanism (regulation of signaling processes, especially receptor desensitization) ^15,35^. An equation was identified that defines this shape, here termed the rise- and-fall to baseline equation (equation (3)). This equation is familiar in pharmacokinetics, being the equation defining drug concentration in the oral dosing absorption and elimination model ^58,59^. We have loaded it into a custom Prism template, “Signaling kinetic model-free equations” available in the supplementary files.

Two examples for second messenger molecules are shown in Fig. 4, diacylglycerol production via the AT_1_ receptor (Fig. 4a) and ^60^cAMP production via the β_2_ adrenoceptor ^49^ (Fig. 4b). In both studies the mechanisms underlying the curve shape were investigated. The rise-and-fall to baseline mechanism was evident when both regulation of signaling mechanisms were operative, i.e. when there were no experimental manipulations of receptor desensitization or degradation of the response. When the regulation mechanisms were manipulated, the shape of the curve changed. When receptor desensitization was blocked or attenuated, the resulting curve shape approached the association exponential curve, as shown in Fig. 2a of the original study for the AT_1_ receptor ^60^ and Fig. 3c for the β_2_ adrenoceptor. Response degradation was in operation, demonstrated for the β_2_ adrenoceptor by the effect of phosphodiesterase inhibition ^49^ and assumed for the AT_1_ receptor because diacylglycerol is typically cleared rapidly by diacylglycerol kinases ^37^.

**Figure 4.**
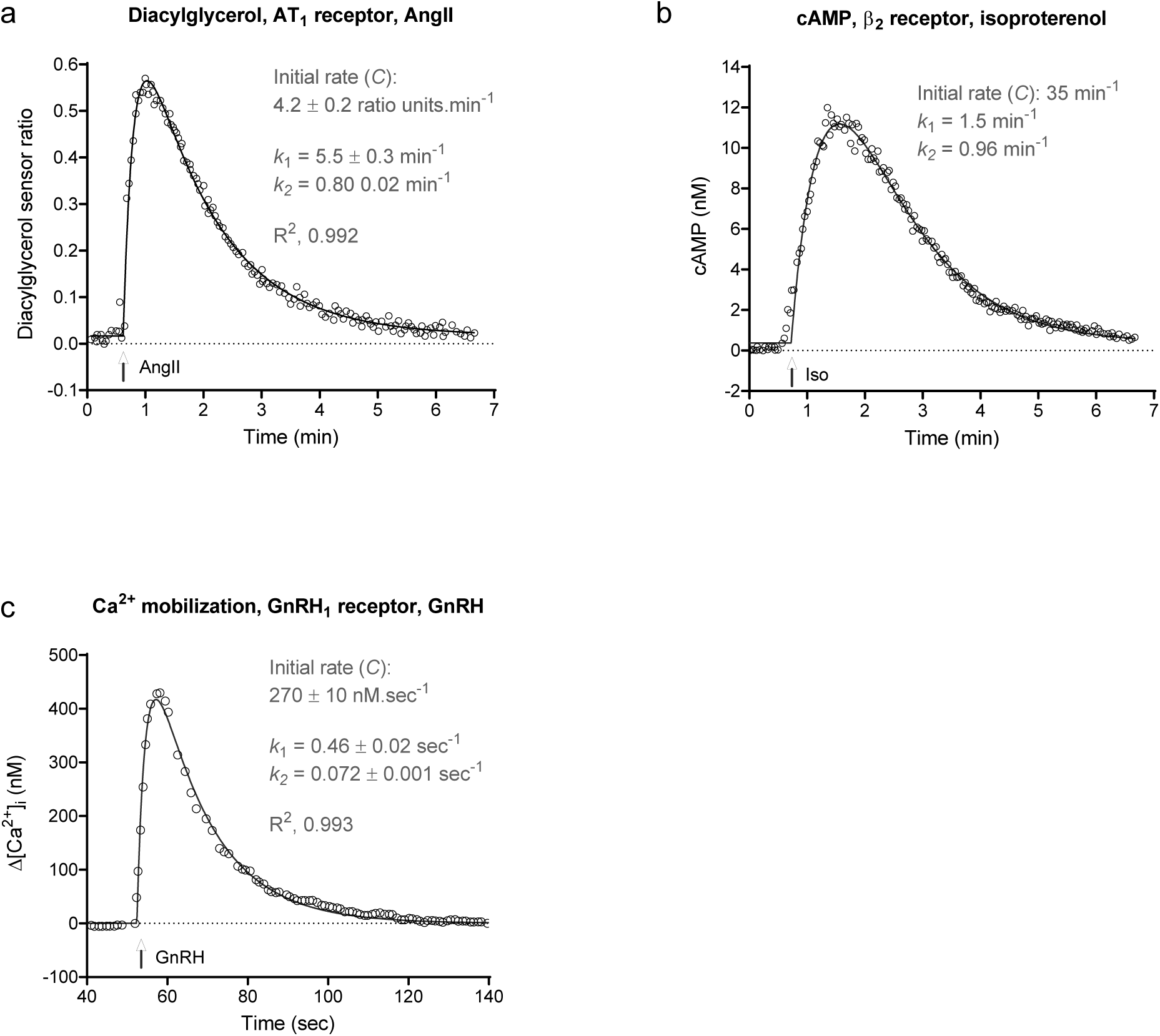
Rise-and-fall to baseline time course profile examples of GPCR signaling. In all cases, none of the regulation of signaling mechanisms were manipulated and the model underlying the curve shape can be inferred. Data were fit to equation (7) in all cases. (a) Diacylglycerol production via the AT_1_ angiotensin receptor stimulated by 100 nM AngII (data from Fig. 1c of ^56^). (b) cAMP production via the β_2_ adrenoceptor stimulated by 10 μM isoproterenol (data from Fig. 3a of ^45^). (c) Intracellular Ca^2+^ mobilization via the GnRH_1_ receptor stimulated by 100 nM GnRH in the absence of extracellular Ca^2+^ (data from Fig. 1a, absence of extracellular Ca^2+^, of ^61^). Note in (b) the fitting procedure involved minimization of outlier contribution to the fit (“Robust regression”) ^70^, which precluded estimation of the fitted parameter error and R^2^ values (see Supplementary file “Curve fit results” for details). The initial rate, equal to *k*_*τ*_, is the fitted parameter *C*.

The rise-and-fall to baseline response is also well known in calcium signaling, representing the change of cytoplasmic Ca^2+^ concentration on stimulation by GPCR agonists (usually when there is no extracellular Ca^2+^ in the assay). An unusual regulation mechanism is involved in Ca^2+^ signaling via intracellular stores. The amount of cytoplasmic Ca^2+^ that can be obtained is limited by the amount of Ca^2+^ in the intracellular stores ^61,62^. Depletion of Ca^2+^ in the store as it is being released into the cytoplasm limits the amount and rate of cytoplasmic Ca^2+^ release. This regulation mechanism can be described as depletion of response precursor ^32^. The response is also regulated by clearance of cytoplasmic Ca^2+^ out of the cell ^63,64^. This can be described as response degradation. These two regulation processes result in the rise-and-fall to baseline profile as described previously ^32^. An example is provided by GnRH-stimulated Ca^2+^ mobilization in pituitary gonadotrophs, as shown in Fig. 4c ^65^. Other receptors giving this profile include the BB1 receptor ^66^ and AT_1_ receptor ^26^. It is important to note that other time course shapes are observed for cytoplasmic Ca^2+^ mobilization, including the rise-and-fall to steady-state curve (considered separately below), and calcium oscillations and waves ^67,68^. In addition, rise-and-fall profiles are observed in Ca^2+^ responses which are not fit well by the rise-and-fall equations used in this study (e.g. in Supplementary Fig. S3).

### Rise-and-fall to steady-state time course profile

A second rise-and-fall curve shape is encountered in GPCR signaling in which the response, after rising to the peak, falls to a plateau level of response which is above baseline, i.e. response declines to a steady-state after peaking. Examples of this shape are shown in Fig. 5. In this study, an equation was identified that describes these data, termed the rise-and-fall to steady-state equation (equation (4)). This equation was identified as a general form of explicit GPCR signaling mechanism equations, as described in Appendix 2. These mechanisms are more complex than the simplest mechanisms considered previously ^32^ and include recently-discovered mechanisms, such as persistent signaling by internalized receptors ^16,17^. These mechanisms are considered below (“More complex models”). We have loaded the equation into a custom Prism template, “Signaling kinetic model-free equations” available in the supplementary files.

**Figure 5.**
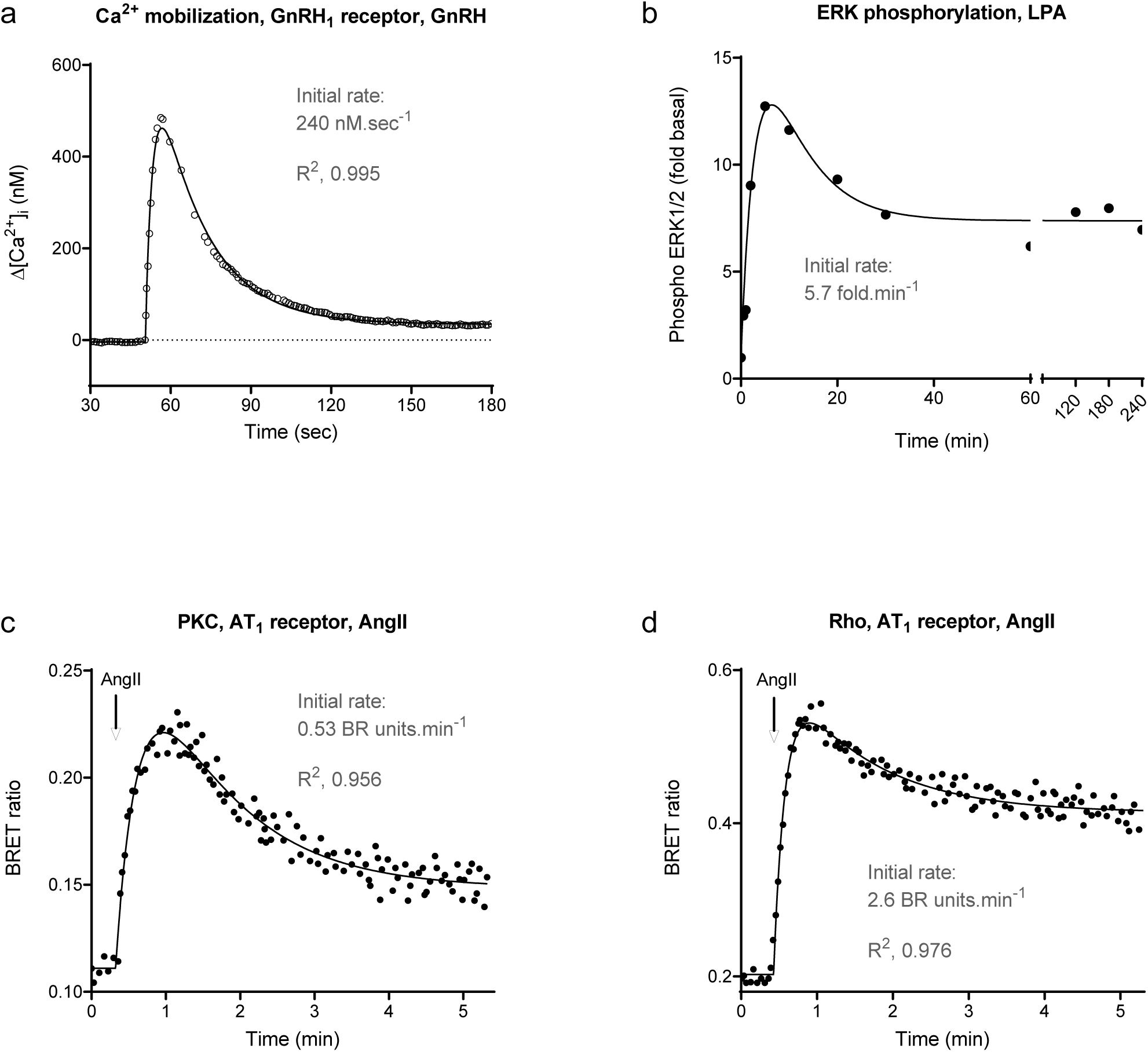
Rise-and-fall to steady-state time course profile examples of GPCR signaling. (a) Intracellular Ca^2+^ mobilization via the GnRH_1_ receptor stimulated by 100 nM GnRH in the presence of extracellular Ca^2+^, an example of the precursor depletion with response degradation to steady-state model (data from Fig. 1a, presence of extracellular Ca^2+^, of ^61^). (b) ERK1/2 phosphorylation via the LPA_1_ receptor stimulated by 10 μM LPA (data from Fig. 4a of ^66^, fit to equation (4) with baseline fixed at 1.0) (c) PKC activation and (d) Rho activation by the AT_1_ receptor in vascular smooth muscle cells stimulated by 100 nM AngII (data from Fig. 3a and b, respectively, of ^67^, fit to equation (8))). Note in (b) the fitting procedure involved minimization of outlier contribution to the fit (“Robust regression”) ^70^, which precluded estimation of R^2^ values. Fitted parameter values are given in the file “Curve fit results” in Supporting Material. Initial rate was calculated using the equation, Intial rate = SSR × (*Dk*_1_ − (*D* − 1)*k*_2_) (See Appendix 1.1). Data in (a) were also analyzed using the mechanism equation for precursor depletion and response degradation to steady-state (equation (17) in Methods). The fitted curve overlies that of the rise-and-fall to steady-state fit.

The most commonly-encountered example was calcium mobilization, the increase of cytoplasmic Ca^2+^ upon application of the GPCR agonist. A representative example is shown in Fig. 5a, cytoplasmic Ca^2+^ concentration stimulated by GnRH via the GnRH_1_ receptor ^65^. It is well known that the cytoplasmic Ca^2+^ level usually reaches a plateau after sustained application of the agonist, particularly when Ca^2+^ is included in the extracellular medium. Under this condition, Ca^2+^ can re-enter the cell and this can result in a steady-state being reached between the numerous processes controlling cytoplasmic Ca^2+^ concentration ^63,67-69^. This mechanism can be represented in the context of the regulation of signaling mechanisms as a reformation of the response from the response decay product, in this case reappearance of cytoplasmic Ca^2+^ from the extracellular space. This mechanism was formulated in Appendix 2.3.3. This mechanism is described in general form by the rise-and-fall to steady-state equation (equation (3)) and in explicit form by equation (15). Data are analyzed by the general form (see “Measuring the initial rate from curve fit parameters” below) and explicit form (“Measuring model parameters”).

The rise-and-fall to steady-state profile is also commonly observed in numerous downstream signaling responses, examples of which are shown in Fig. 5: ERK1/2 phosphorylation by lysophosphatidic acid ^70^, and protein kinase C and Rho activation via the AT_1_ receptor ^71^. In some cases, the data fit the rise-and-fall to steady-state equation well (high R^2^ value) but the fitted parameters were ambiguous, a scenario encountered when the two rate constant values were almost equal. An example is provided in Supplementary Fig. S4. This observation requires further investigation, for example using structural identifiability analysis.

### Initial rate measurement using model-free analysis

In order for time course analysis to be useful for pharmacological investigations, pharmacological parameters need to be extracted from the data that capture the temporal dimension of activity. One kinetic parameter used routinely for another target class (enzymes) and occasionally for GPCRs is the initial rate of activity. This is the rate at the earliest part of the time course, when the response increases linearly over time. This parameter offers several benefits. 1) It is a biologically-meaningful descriptor of signaling activity. 2) In principle, the initial rate is effectively a pure efficacy parameter, being the rate of signal generation before it is affected by regulation of signaling mechanisms, which affect the shape of the later part of the time course. This is formally demonstrated using the kinetic pharmacological model below. 3) From a practical perspective, the initial rate provides a single parameter, as opposed to multiple parameters, of drug efficacy, simplifying application to ligand optimization in medicinal chemistry projects. 4) The initial rate can be determined regardless of the shape of the time course, enabling comparison of a ligand’s efficacy between responses with different temporal profiles (see below). We recently applied the initial rate approach to arrestin-receptor interaction, a special case in which there was no signal transduction ^26^. Here it is applied universally to GPCR signaling responses.

### Measuring the initial rate from curve fit parameters

For curved time course data, the initial rate is often measured by assessing which data points lie on the linear portion at the start of the curve, then fitting a straight line equation to those points. Here a more efficient method is developed. The entire time course is fit to the equation that defines the curve, then the initial rate calculated from the fitted parameters. This requires an additional equation that defines the initial rate (IR) in terms of the fitted parameters. This equation was obtained here for each of the time course shapes, as the limit of the time course equation as time approaches zero (the formal definition of the initial rate condition) (Appendix 1):

Straight line, from equation (1):

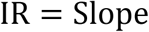

Association exponential, from equation (2):

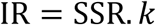

Rise-and-fall to baseline, from equation (3):

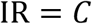

Rise-and-fall to steady-state, from equation (4):

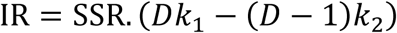

This method was used to quantify the initial rate of signaling for the responses in Figs. 2-5 and Supplementary Figs. S1,S2. Data were fit to the appropriate time course equation using Prism 8. For linear and association exponential profiles data were analyzed using built-in equations provided with the software. For the rise-and-fall profiles, user-entered custom equations were used. (A ready-to-use Prism template containing these equations, “Signaling kinetic model-free equations,” is provided in the supplementary files.) In some cases, the signal initiation time was a variable in the equation (see equations 5-9), allowing for delay between addition of ligand and initiation of signal (e.g. in gene expression assays, Supplementary Fig. S1), or accommodating the scenario where the ligand addition time is not precisely defined. The data presented in this study were fit well by the equations; in all but one case, the correlation coefficient was > 0.95, and the standard error of the fit parameters was typically < 10% of the fitted value (see “Curve fit results” Excel file in Supporting Material). With two exceptions, data were fit well using the default fitting method of Prism 8 (least squares regression with medium convergence criteria ^72^). The exceptions were the rise-and-fall fit in Fig. 4b (cAMP production via the β2 adrenoceptor without phosphodiesterase inhibition) and Fig. 5b (LPA-stimulated ERK phosphorylation). In these cases, the default method yielded an ambiguous fit ^73^ (see “Curve fit results” Excel file in Supporting Material), which was resolved using the “Robust regression” option which minimizes the contribution of outliers to the fit ^74^.

The initial rate value was then calculated from the fitted parameter estimates using the equations above. Note the simplicity of obtaining the initial rate in most cases. For example, for the association exponential curve, the initial rate is simply the rate constant multiplied by the steady-state response. For the rise-and-fall to baseline curve, it is simply the value of the fitted parameter *C*. The calculated initial rate values are shown in the figures (Figs. 2-5, Supplementary Figs. S1,S2). A maximally-stimulating concentration of agonist was employed in all the examples provided. Under this condition, the initial rate is the efficacy parameter of the agonist in kinetic terms, here termed IR_max_.

### Application to biased agonism of the V_2_ vasopressin receptor

The model-free method was applied here to G-protein signaling and arrestin recruitment by the V_2_ vasopressin receptor. This receptor is activated by two cognate endogenous ligands, vasopressin and oxytocin ^75^. The receptor is well known to couple stably to arrestin when bound by vasopressin (a so-called class B arrestin binding profile ^33^) and provides an example of biased agonism by endogenous ligands; oxytocin is biased toward G-protein over arrestin, relative to vasopressin ^76^.

Here the model-free method was used to quantify signaling and evaluate biased agonism in kinetic terms, by quantifying the initial rate of the responses (cAMP accumulation and arrestin-3 recruitment). Genetically-encoded biosensors were used to quantify these responses in real time in live cells expressing the receptor ^26,77^, as described in Methods. The concentration-dependence was determined, using a range of concentrations of vasopressin or oxytocin. The time course profiles are shown in Fig. 6. In all cases, the data were fit well by the association exponential equation (equation (6), Fig. 6a,b,de, see also “Curve fit results” Excel file in Supporting Material). From the fitted parameters, the initial rate was calculated as the steady-state response multiplied by the rate constant (“Curve fit results” Excel file). This was done for all concentrations of the two ligands for the two pathways. The concentration-response of the initial rate was then evaluated, as shown in Fig. 6c,f. The concentration response was fit well by a standard sigmoid curve equation, from which the efficacy and potency of the ligand could be determined. The efficacy is the maximal initial rate (IR_max_), i.e. the upper plateau of the curve. The potency is the midpoint concentration of the ligand, specifically the concentration of ligand giving an initial rate half of the maximum value.

**Figure 6.**
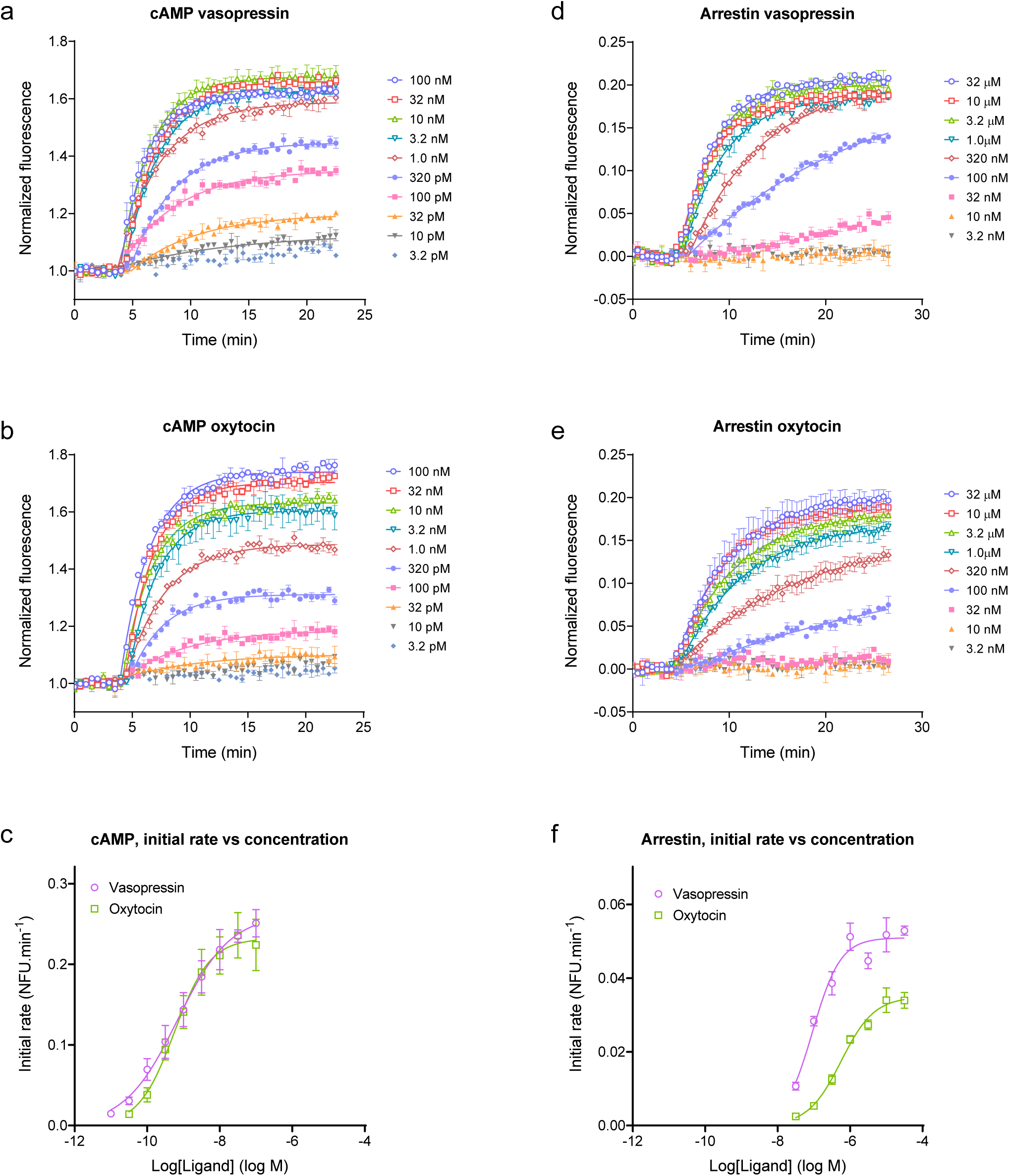
Applying the model free initial rate analysis to V2 vasopressin receptor-mediated cAMP signaling and arrestin recruitment. The time course of cAMP production (a,b) and arrestin recruitment (d,e) were measured using genetically-encoded biosensors ^22,73^, as described in Methods, for vasopressin and oxytocin. Time course data were well fit by the association exponential equation (equation (6), with fitted signal initiation time, fitted values in “Curve fit results” Excel file in Supporting Material). The initial rate at each concentration was calculated by multiplying the steady-state response by the rate constant value (SSR × *k*). The initial rate was then plotted versus the ligand concentration (c,f) and the data fitted to a sigmoid curve equation ^95^, providing fitted values of the maximal initial rate (IR_max_, equal to *k*_*τ*_) and the L50 (concentration of agonist producing 50% of the maximal initial rate). The fitted values are given in Table 1. Data points are mean of two or four technical replicates from representative experiments in the time course (a,b,d,e). Error bars are SEM. Data points in the concentration-response graphs (c,f) are mean ± SEM of the initial rate values from three independent experiments.

The concentration-response analysis indicated oxytocin was less active for recruiting arrestin than vasopressin (Fig. 6f, Table 1), consistent with a previous report ^76^. This was manifest as a lower maximal initial rate of arrestin recruitment (IR_max_ value 70 % of the vasopressin value) and also as a 6.2-fold lower potency (580 nM vs 93 nM for vasopressin). These results indicate the weaker arrestin recruitment by oxytocin can be accounted for by a slower initial rate of recruitment by the oxytocin-bound receptor, and a lower affinity of oxytocin compared with vasopressin for the arrestin-bound receptor. By contrast the two ligands were effectively equivalent for stimulation of cAMP production (Fig. 6c, Table 1). These findings indicate biased agonism of oxytocin relative to vasopressin (G-protein bias) and are consistent with oxytocin binding / stabilizing a different conformation of the V_2_ receptor compared with that bound by vasopressin.

**Table 1.** V_2_ initial rate dose response for cAMP generation and arrestin recruitment stimulated by vasopressin and oxytocin.

| Ligand | cAMP |  |  | Arrestin |  |  |
| --- | --- | --- | --- | --- | --- | --- |
| | $IR_{max}, k_{\tau}$<br>NFU.min<sup>-1</sup> | $IR_{max}, k_{\tau}$<br>% vaso. | - log L<sub>50</sub><br>(L<sub>50</sub>, nM) | $IR_{max}, k_{\tau}$<br>NFU.min<sup>-1</sup> | $IR_{max}, k_{\tau}$<br>% vaso. | - log L<sub>50</sub><br>(L<sub>50</sub>, nM) |
| Vasopressin | 0.26 ± 0.01 | 100 ± 0 | 9.16 ± 0.20<br>(0.70) | 0.051 ± 0.003 | 100 ± 0 | 7.03 ± 0.06<br>(93) |
| Oxytocin | 0.24 ± 0.05 | 90 ± 8 | 9.21 ± 0.18<br>(0.61) | 0.035 ± 0.003 | 70 ± 9 | 6.23 ± 0.06<br>(580) |

### Mechanistic pharmacological model of GPCR signaling and regulation kinetics

Recently a mechanistic kinetic pharmacological model was developed to quantify GPCR signaling activity in kinetic terms ^32^ that is being applied to understanding signaling efficacy in kinetic terms ^8^. The model comprises a minimal number of parameters to enable parameter estimation by curve fitting. The approach is macroscopic – ligand signaling activity is defined at the level of the whole system, like the operational model of agonism ^7^. (Numerous microscopic systems-biology type models have been developed to simulate GPCR signaling dynamics that quantify each interaction in the signaling cascade ^8,49,78-82^ but, while they enable close examination of mechanism, they contain too many parameters to be useful in medicinal chemistry campaigns.) The model is illustrated in Fig. 7. Signaling activity is quantified using a single, readily-measurable parameter, *k*_*τ*_, which is the initial rate of signaling by the agonist-bound receptor in enzymatic terms ^32^. The model is extended to incorporate known regulation of signaling mechanisms. GPCR signaling is regulated to limit signaling, preventing over-stimulation of the cell. We consider the properties of the model, how it relates to the model-free analysis, and how to estimate the efficacy and regulation parameters by curve fitting. The model gives rise to the four curve shapes observed experimentally. It emerged that the specific shape is simply dependent on the number of regulatory mechanisms (0, straight line; 1, association exponential; 2, rise-and-fall to baseline). It is shown the model-free analysis emerges from the mechanistic model, the equations of the former being general forms of the equations for the latter. Estimating efficacy is straightforward and does not require knowledge of the mechanism - it is shown *k*_*τ*_ is equal to IR_max_. The regulation parameters can be estimated if the mechanism is known, for example the receptor desensitization rate.

**Figure 7.**
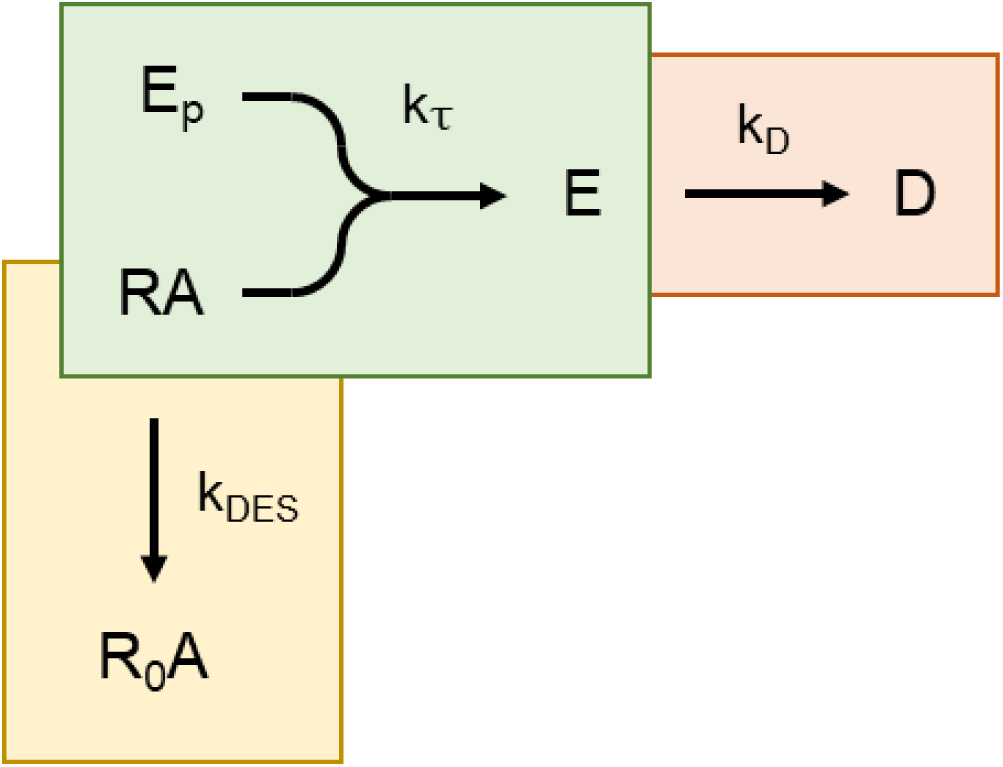
Kinetic mechanistic model of agonism incorporating regulation of signaling. Both mechanisms of short-term signaling regulation are included – receptor desensitization (yellow) and response degradation (pink). The response generation process is in green. In this process, the response precursor (*E*_*P*_) is converted to the response (*E*) by the agonist-occupied receptor (*RA*). This proceeds at a rate defined by *k*_*τ*_, the transduction rate constant, which is the initial rate of response generation by the agonist-occupied receptor. Receptor desensitization is represented by transformation of *RA* into the inactive receptor *R*_0_*A* that does not generate a response (because it can’t couple to *E*_*P*_). The rate constant for desensitization is *k*_*DES*_. Degradation of response is represented by decay of *E* to *D*, governed by the degradation rate constant *k*_*D*_. This model is an extension the original kinetic mechanistic model ^28^, extended here to incorporate receptor desensitization.

### Model description

The kinetic mechanistic model describes GPCR signaling in terms of enzyme activity. A signal results from conversion of a signal precursor (analogous to the substrate) to the signal (the product) by the agonist-bound GPCR (the enzyme) (Fig. 7), as described previously ^32^. The rate of ligand efficacy for generating the response is quantified as the initial rate of signaling, termed *k*_*τ*_, analogous to the initial rate of enzyme activity. *k*_*τ*_ is the rate of response generation by the agonist-occupied receptor, defined as the product of the total receptor concentration ([*R*]_*TOT*_), total signal precursor (*E*_*P*_) and the response-generation rate constant (*k*_*E*_):

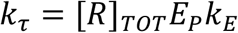

(By comparison, the initial rate of enzyme activity is the product of the enzyme concentration, substrate concentration and the catalytic rate constant.) The model as formulated in this study assumes receptor-ligand occupancy does not change over time, that the equilibrium level of occupancy is rapidly achieved and is not rate-liming. This condition is likely met for maximally-stimulating ligand concentrations (used to quantify efficacy), and for all concentrations of lower potency ligands, scenarios which result in rapid association of ligand with receptor (see Discussion). Note the model framework is amenable to extension to incorporate the kinetics of receptor-ligand binding, as described previously ^32^.

The model can be extended to incorporate regulation of signaling. The canonical mechanisms are response degradation and receptor desensitization. Response degradation is the process by which the signal, once generated, is cleared over time. Examples include breakdown of second messenger molecules, such as cAMP by phosphodiesterases ^36^, and de-activation of G-protein by hydrolysis of bound GTP to GDP by the intrinsic GTPase activity of the G-protein ^83^. Some signals are decreased by clearance of the signaling species from the relevant compartment, for example efflux of cytosolic Ca^2+^ ions ^64^. In the model, response degradation (pink region of Fig. 7) is represented simply by exponential decay, governed by *k*_*D*_, the response degradation rate constant. This component of the model was described previously ^32^.

### Receptor desensitization model

In the present study, the model is extended to incorporate the second canonical regulation process, receptor desensitization (Fig. 7 yellow region). Receptor desensitization typically results from receptor phosphorylation and subsequent arrestin binding, which inhibits G-protein interaction ^33,35^. This process is represented simply as an exponential decay of the agonist-bound receptor concentration that can couple to the signaling machinery. This is governed by the desensitization rate constant, *k*_*DES*_. The basic model is formulated in Appendix 2.1 and is represented by Scheme 1 below:

**Scheme 1.**
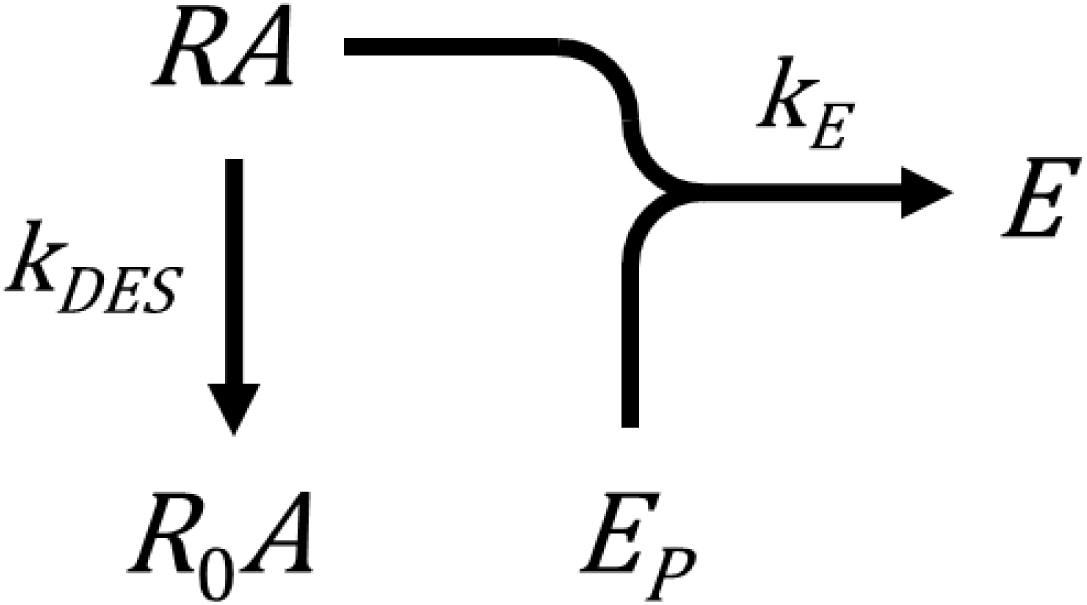

The time course shape predicted by this model is an association exponential curve (Fig. 8b). This is consistent with experimental results; systems in which the sole regulation mechanism is receptor desensitization yield an association exponential time course. Examples include IP production by the AT_1_ (Fig. 3a)^39^, PAR1 (see Fig. 2b in ^40^) and C-terminally-extended GnRH receptor (see Fig. 3c in ^42^) (when response degradation is blocked by Li^+^). This curve shape makes sense intuitively. At early times response is generated rapidly. The response then slows, because response generation is attenuated by the loss of active receptor. Ultimately the response approaches a limit (the plateau). At this limit the response level does not change because no new response is being generated and the existing response is not degraded.

**Figure 8.**
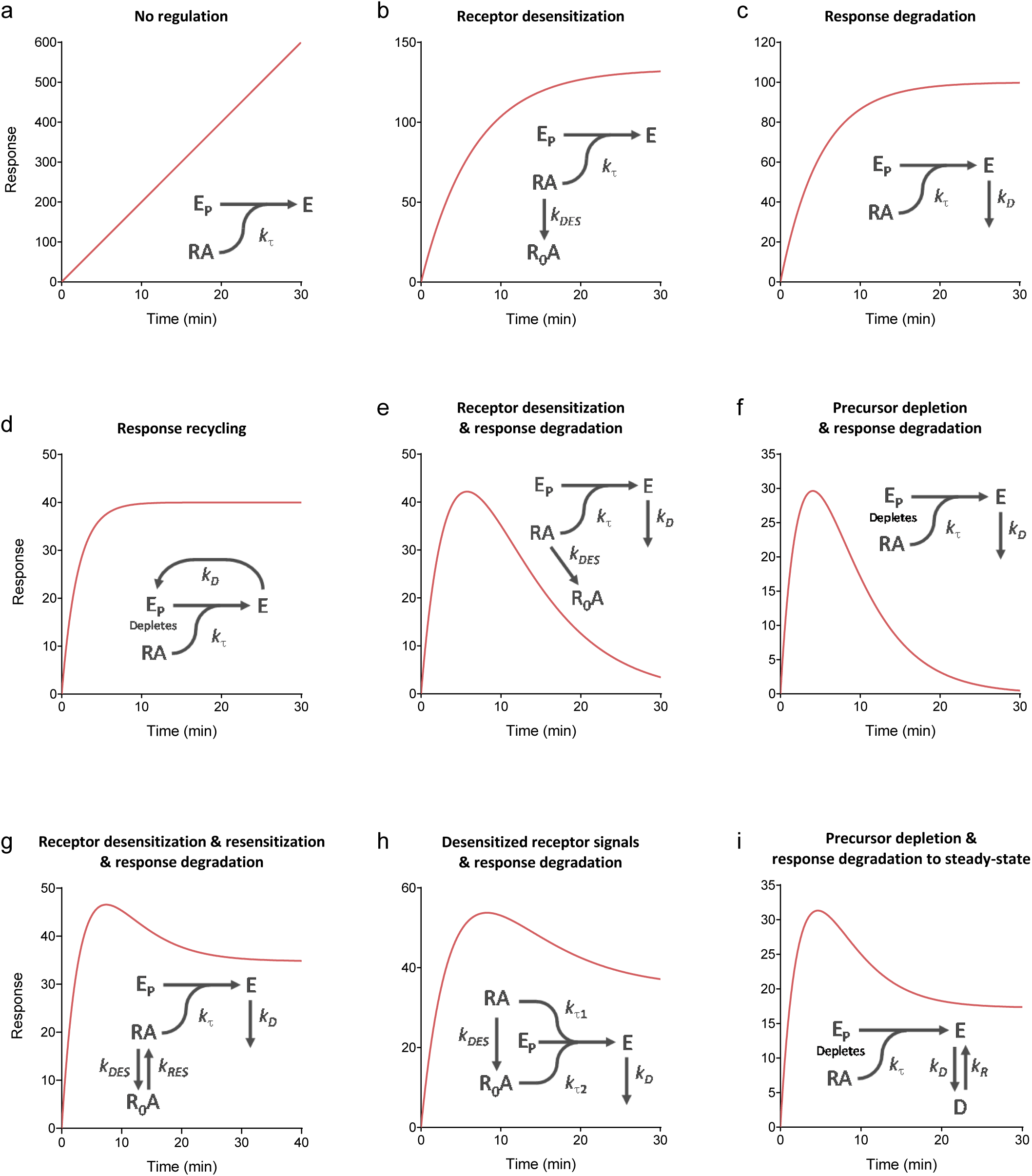
Kinetic signaling and regulation model time course simulations. Data were simulated using the model equations in Supplementary Tables S1-S4, with terms defined in Supplementary Table S5. Parameter values were: *k*_*τ*_, 20 response units; *k*_*DED*_, 0.15 min^-1^; *k*_*D*_, 0.2 min^-1^; *k*_*DED*_, 0.3 min^-1^; *k*_*RED*_, 0.08 min^-1^; *k*_*τ*1_, 20 response units; *k*_*τ*2_, 7 response units; *k*_*R*_, 0.07 min^-1^.

This model can now be extended to incorporate both regulation mechanisms, receptor desensitization and response degradation (Appendix 2.2), represented by Scheme 2 below:

**Scheme 2.**
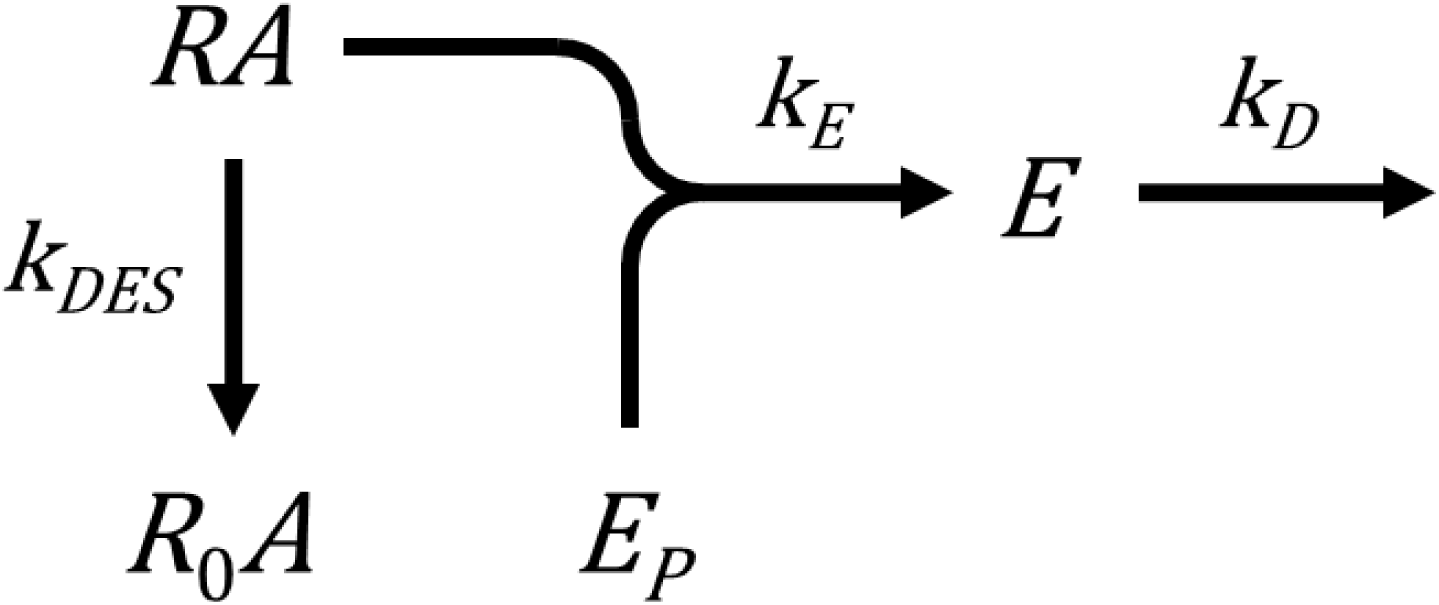

The time course shape for this model is a rise-and-fall to baseline curve (Fig. 8e). This makes sense intuitively. The response rises rapidly then slows as receptor desensitization starts to slow the rate of response generation. The response becomes further limited owing to response degradation. The response reaches an upper limit (the peak) when the rate of response generation equals the rate of degradation. After this, response degradation predominates over response generation. Less and less new response is generated because the active receptor concentration is declining, and ultimately no new response will be generated because the active receptor concentration will decline to zero. Ultimately the response level falls to zero once all existing response has been degraded. This profile is evident in systems where both mechanisms have been shown to be in operation. Examples include diacylglycerol production via the AT_1_ receptor (Fig. 4a) ^60^, cAMP accumulation via the β_2_ adrenoceptor in the absence of phosphodiesterase inhibition (Fig. 4b) ^49^, and IP production via the CCK1 receptor in the absence of Li^+^ (Fig. 10a of ^84^).

### More complex models

More complex regulation and signaling mechanisms discovered experimentally can be represented using the model formulation, as described in Appendix 2.3. Receptor can resensitize and then contribute to signaling. This model is formulated in combination with response degradation in Appendix 2.3.1. The receptor, once desensitized and internalized, can continue to signal, as first described for parathyroid hormone-1 and sphingosine 1-phosphate-1 receptors ^16,17^. This model is formulated in Appendix 2.3.2. A third more complex model describes calcium signaling; response is degraded (cleared from the cytoplasm) but the response can reform (from calcium that re-enters the cell). This process, in combination with depletion of the response precursor, is formulated in Appendix 2.3.3. In all three models the time course is described by a rise-and-fall to steady-state equation (see Appendix 2.3).

### Comparison of models with experimental data

The receptor desensitization model completes the set of kinetic mechanistic signaling models developed for GPCRs, initiated in ref. ^32^. This allows comparison of a complete kinetic model of GPCR signaling and regulation with experimental data, and with the model-free analysis. The model mechanisms are shown schematically in Fig. 8 and are: no regulation; receptor desensitization; response degradation; response recycling; receptor desensitization and response degradation; precursor depletion and response degradation; and the three more complicated models described above and in Appendix 2.3. Simulators of the models are provided in Supporting Material, enabling investigators to evaluate changes of the parameter values.

The time course curve shapes predicted by the model are those observed experimentally – the straight line, the association exponential, and the two rise-and-fall curves (Fig. 8). It emerges that the shape is dependent on the number of regulatory mechanisms. With no regulatory mechanisms, the time course is a straight line (Fig. 8a). With one mechanism, an association exponential curve results (receptor desensitization, response degradation, or response recycling, Fig. 8b-d). With two mechanisms, a rise-and-fall to baseline curve results (Fig. 8e,f). More precisely, when an input regulation mechanism (desensitization or precursor depletion) is coupled with an output mechanism (response degradation). With the more complicated mechanisms, a rise-and-fall to baseline profile results (Fig. 8g-i).

### Comparison of models with model-free analysis

The four curve shapes of the mechanistic model are the same as those of the model-free analysis. This is shown graphically by comparing Fig. 8 with Fig. 1. It is also evident mathematically by inspection of the equations; the model equations listed in Supplementary Tables S1-S4 conform to the forms of the general equations (equations (1-4)). This finding indicates the model-free analysis emerges from known mechanisms of GPCR signaling and regulation.

The maximal initial rate from the model-free analysis (IR_max_) is equal to *k*_*τ*_ from the mechanistic model. This is evident by taking the initial rate of the equations, i.e. the limit as time approaches zero. By definition this limit is IR_max_ for the model-free analysis. For the mechanistic equations, this limit is *k*_*τ*_ (for a maximally-stimulating concentration of agonist, Appendix 1.2). This supports the hypothesis that the initial rate is purely an efficacy term, being the rate of response generation before it is impacted by regulation of signaling mechanisms, because *k*_*τ*_ contains efficacy terms but no regulation terms. It also provides a mechanistic foundation of IR_max_, indicating it is analogous to the initial of an enzyme’s activity (formally, the product of the receptor and precursor concentrations and a rate constant).

### Measuring model parameters

The models contain parameters for signal generation and for regulation of signaling. These parameters can be estimated by curve fitting by two methods. The time course data can be fit directly to equations that explicitly describe the model. These equations are listed in Supplementary Information and are provided in a custom Prism template in the supplementary files (“Signaling kinetic mechanism equations”). Alternatively, the data can be fit to general time course equations (equations (1-4)) and the fitted parameter values used to calculate the model parameters. These approaches are used here to estimate efficacy and regulation parameters, which can be done using just a maximally-stimulating concentration of agonist. Note in all the literature examples in this study, a maximally-stimulating concentration was used.

The general equation fitting approach is used here for the linear, association exponential, and rise-and-fall to baseline examples. For the linear examples, the only parameter to be estimated is *k*_*τ*_ and this is equal to the slope of the line for a maximally-stimulating concentration of agonist. The resulting *k*_*τ*_ values are the Slope values in Fig. 2 and Supplementary Fig. 1.

For the association exponential time course, there are two parameters, the efficacy parameter *k*_*τ*_, and a regulation parameter specified by the model (for example, the desensitization rate constant *k*_*DES*_). *k*_*τ*_ can be estimated using the same method as that used in the model-free analysis to calculate IR_max_, since these parameters are equivalent (see “Measuring the initial rate from curve fit parameters” above). The rate constant *k* is multiplied by the steady-state level of response, for a maximally-stimulating concentration of agonist. This resulting *k*_*τ*_ values are equal to the initial rate values given in Fig. 3 and Supplementary Fig. 2, and the IR_max_ values in Table 1. Note in all the model variants that give rise to the association exponential profile, *k*_*τ*_ is the product of *k* and the steady-state response, i.e. *k*_*τ*_ is calculated in the same way. This means *k*_*τ*_ can be estimated without knowing the specific mechanism underlying the curve shape. The regulation parameter of the association exponential time course is dependent on the mechanism. In all cases, the regulation parameter is given by the rate constant parameter *k* of the general equation (Supplementary Table S2). If the mechanism is known, the value of *k* can be ascribed to a regulatory process. This is shown using the studies cited here where the regulation of signaling mechanism was evaluated. In Fig. 3a, IP generation by the AT_1_ receptor ^39^, the mechanism was most likely receptor desensitization since deletion of arrestin changes the shape to a straight line (Fig. 2a) and response degradation was blocked by Li^+^. The rate constant *k* then represents the desensitization rate constant *k*_*DES*_. The fitted value of 0.040 min^-1^ indicated a receptor desensitization half time of 17 min. In Fig. 3b and c, the mechanism is most likely response degradation since receptor desensitization was minimized, the GnRH receptor lacking a C-terminal tail ^44^ (Fig. 3b), the β2 adrenoceptor-expressing cells devoid of arrestin ^39^ (Fig. 3c), and in both cases inhibitors of degradation excluded. The *k* value then represents the response degradation rate constant *k*_*D*_. The fitted values, 0.12 min^-1^ for the GnRH_1_ receptor and 1.1 min^-1^ for the β2 adrenoceptor, indicated response degradation half-times of 5.8 min and 38 sec, respectively.

For the rise-and-fall to baseline mechanisms, there are three parameters – the efficacy parameter *k*_*τ*_, and two regulation parameters. *k*_*τ*_ can again be estimated using the general equation without knowing the specific mechanism underlying the time course profile. *k*_*τ*_ is equal to the parameter *C* of the general equation (equation (3)) when a maximally-stimulating agonist concentration is employed, and these values are shown for the literature examples in Fig. 4. The rate constant values can be ascribed to the mechanism rate constants if the mechanism is known. In Fig. 4a, diacyglycerol production by the AT_1_ receptor, the mechanism is most likely receptor desensitization and response degradation, since blocking arrestin recruitment changes the shape to an association exponential (see Fig. 2a of ref. ^60^) and diacyclglycerol is rapidly degraded by diacylglycerol kinases ^37^. It is then necessary to determine which of the general rate constants (*k*_1_ and *k*_2_) corresponds to which of the mechanism rate constants (*k*_*DES*_ and *k*_*D*_). In this case, it is likely the faster of the rates (*k*_1_) corresponds to *k*_*D*_, since blocking arrestin recruitment gives an association exponential profile with rate similar to *k*_1_ (see Fig. 2a of ref. ^60^). This approach gives a *k*_*D*_ (*k*_1_) value of 5.5 min^-1^ and a *k*_*DES*_ (*k*_2_) value of 0.80 min^-1^, corresponding to degradation and desensitization half times of 7.6 sec and 52 sec, respectively. In Fig. 4c, cytoplasmic Ca^2+^ elevation via the GnRH receptor, the mechanism is most likely precursor depletion and response degradation, based on the known mechanism of calcium responses (see “Rise-and-fall to baseline time course profile” above). The more rapid of the two rate constants is most likely precursor depletion since this is the first event in the signaling cascade. This logic gives a *k*_*DEP*_ (*k*_1_) value of 0.46 sec^-1^ and a *k*_*D*_ (*k*_2_) value of 0.072 sec^-1^, corresponding to depletion and degradation half times of 1.5 sec and 9.6 sec, respectively. Fig. 4b shows cAMP generation by the β2 adreneoceptor. In this example, likely resulting from receptor desensitization and response degradation based on the experiments in ref. ^49^, the two rate constant values are close to one another (1.5 and 0.96 min^-1^), preventing the ascribing of the constants to the regulatory processes but demonstrating the rate of the processes is similar in this system.

For the rise-and-fall to steady-state models, *k*_*τ*_ can be estimated using the general equation and a maximally-stimulating concentration of agonist. *k*_*τ*_ is equal to IR_max_, calculated from the parameters as described above. For estimating the regulation parameters, we recommend using the mechanistic model equation rather than the general equation. For this curve shape, the general equation parameters correspond to complicated combinations of the mechanism parameters, which introduces constraints of the parameter values that are not incorporated into the general model equations (Appendix 2.3). This approach was used to analyze the Ca^2+^ mobilization data in Fig. 5a. In this experiment, Ca^2+^ was included in the extracellular medium, which results in a steady-state being reached between cytoplasmic Ca^2+^ entry and export. This is the “Response degradation to steady-state with precursor depletion” model (Appendix 2.3.3). The equation fitted the data well (R^2^ value 0.995, see curve in Fig. 5a, standard error of the fitted model parameters less than 10% - see “Curve fit results” Excel file in Supporting Material). The regulation rate constant values were 0.37 sec^-1^ for *k*_*DEP*_, 0.052 sec^-1^ for *k*_*D*_ and 0.0033 sec^-1^ for *k*_*R*_, corresponding to half times for precursor depletion, response degradation and response reformation of 1.9 sec, 13 sec and 3.5 min, respectively. As expected, the *k*_*τ*_ value from the mechanism equation fit was the same as the value from the model-free analysis (240 nM.sec^-1^).

## Discussion

New technologies have enabled efficient measurement of the kinetics of GPCR signaling using continuous-read, real-time bioassay modalities ^12-14^. Methods are now required to extract pharmacological parameters from the time course data. Such methods have been developed for specific responses that are immediately proximal to the receptor (internalization ^25^ and arrestin-receptor interaction ^26^). Here methods were designed for universal application to GPCR signaling and regulation responses. Curve-fitting methods are described for analyzing time course signaling data, designed for routine use by pharmacologists, enabling application to medicinal chemistry and receptor research. The equations are simple and built into commonly-used curve-fitting programs, or are provided in a ready-to-use templates. The efficacy for signaling is quantified kinetically by an intuitive, familiar and biologically meaningful parameter, the initial rate of signaling, which is obtained from the curve fit parameters. The underlying theoretical framework is familiar, based on the concepts of enzyme kinetics and the known mechanisms of regulation of signaling. Resources are provided in Supporting Material to facilitate implementation and application of the models, including Prism templates containing the equations, and a time course data simulator for the mechanistic models.

This approach has practical benefits for optimization of new molecules and for GPCR signaling research. In project workflow, it allows representation of the time course data set by a minimal number of informative parameters (e.g. *k*_*τ*_, *k*_*DES*_). This enables tabulation of kinetic signaling data for a series of ligands, necessary for medicinal chemists to establish kinetic structure-activity relationships. If the response at a specific time point is required, for example when comparing in vivo and in vitro activity, this can be calculated from the curve fit parameters, avoiding the necessity of measuring precisely the same time points in every experiment. The approach solves the time-dependency problem, in which pharmacological parameter values can be dependent on the time point at which they are measured, complicating quantification of efficacy and biased agonism ^24-26^. This is because: 1) Activity is quantified as a rate constant value, which is constant over time; 2) Differences of curve shape can be accommodated because the same efficacy parameter, the initial rate, can be extracted from all four of the commonly-encountered time course curve shapes. Finally, the method separates efficacy and regulation, allowing these parameters to be quantified separately (e.g. *k*_*τ*_ and *k*_*DES*_), enabling independent structure-activity optimization of these processes.

The mechanistic model yields insights that explain the time course curve shapes and provide mechanistic meaning to the parameters from the model-free analysis. The model and experimental data show that regulation of signaling defines the time course curve shape. Regulation changes the shape from a straight line (where response is generated indefinitely), to an association exponential curve or a rise-and-fall curve, with the type of curve dependent on the number and nature of the regulation mechanisms (Fig. 7). The underlying mechanism can be interrogated by blocking receptor desensitization and/or response degradation. The model supports the hypothesis that the initial rate of activity is purely an efficacy term, unaffected by regulation of signaling: It is shown that *k*_*τ*_, devoid of regulation terms, is the limit of the equations as time approaches zero, the formal definition of the initial rate (Appendix 1). The model translates the empirical parameters of the model-free analysis into biologically-meaningful parameters of GPCR signaling mechanisms. IR_max_ is equivalent to *k*_*τ*_, the initial rate of signaling generation by the agonist occupied receptor, analogous to the initial rate of an enzyme’s activity. The rate constants *k, k*_1_ and *k*_2_ correspond simply to the rates of the regulation processes, e.g. receptor desensitization and response degradation. If the mechanism is known, the rate constant value can be assigned to a regulatory activity, enabling regulation of signaling to be quantified in simple, kinetic terms.

The relationship between the kinetic analysis and existing pharmacological analysis is now considered. Established pharmacological analysis quantifies signaling in terms of efficacy and affinity, usually at a single time point at which equilibrium between receptor and ligand is assumed ^3-8^. The efficacy term in the kinetic approach (IR_max_ or *k*_*τ*_) is a direct analogue of established efficacy parameters (E_max_, and *τ* of the operational model ^7^). Consequently, the rank order of efficacy should be the same using both approaches. Affinity is more difficult to measure using the kinetic approach because of the equilibration issue. Properly measuring affinity from a concentration-response requires the assay to be incubated long enough for equilibrium between receptor and ligand to be closely approached ^85^. By contrast, the initial rate in the kinetic method is quantified using the earliest time points after the addition of ligand and under these conditions it cannot be generally assumed the ligand and receptor are at equilibrium ^5,86-88^. The impact can be evaluated by considering typical values of receptor-ligand binding rate constants, using the mass-action receptor-ligand association kinetics equation ^89^. For low affinity ligands, the equilibration time is short, potentially enabling accurate estimation of affinity when using responses that proceed over several minutes. A compound with 10 μM affinity binding with an association rate constant (*k*_on_) of 10^8^ M^-1^min^-1^ reaches 97 % of its equilibrium occupancy within 0.1 seconds at its *K*_d_ concentration (i.e. 10 μM). However, for high affinity ligands the equilibration time can be long relative to the duration of the response. A compound with 1 nM affinity and a *k*_on_ of 10^8^ M^-1^min^-1^ takes 18 min to reach 97 % of its equilibrium receptor occupancy when applied at its *K*_d_ concentration. The problem is magnified when the response is very rapid, as previously described ^87^, particularly in Ca^2+^ signaling, which proceeds in seconds. This issue can be circumvented when receptor and ligand can be pre-incubated prior to the initiation of the response, for example in GTPγS binding assays, but this option is rarely available in whole cell assays, the most commonly used assay modality. Importantly, this issue does not affect estimation of efficacy. IR_max_ and *k*_*τ*_ are estimated using maximally-effective concentrations which, due to mass action, bind more rapidly. For example, when applied at 10 μM concentration, the 1 nM *K*_d_, 10^8^ M^-1^min^-1^ *k*_on_ compound reaches 97 % of equilibrium occupancy within 0.2 seconds. Finally, the effect of signal amplification, manifest as receptor reserve, in the kinetic mechanistic model requires further investigation. While the model can incorporate receptor reserve intrinsically (as a depletion of response precursor) ^32^, the rate of signaling in a multistep pathway is ultimately a function of the rates of the individual steps and the stoichiometric relationships between them. For example, the initial rate could be susceptible to receptor reserve effects if there is a rate-limiting step upstream of the response being measured. This issue requires experimental investigation.

In conclusion, this study introduces straightforward data analysis methods to quantify the kinetics of GPCR signaling. The simplicity of the analysis procedures, the intuitive nature of the parameters, and the mechanistic foundation in known and emerging GPCR signaling paradigms will facilitate pharmacological discovery and optimization in kinetic terms.

## Supporting information

Curve fit results

Kinetic mechanism simulations

## Acknowledgements

Research reported in this publication was supported by National Institute of General Medical Sciences, National Institutes of Health under award number R44GM125390 and National Institute of Neurologic Disorders and Stroke, National Institutes of Health R44NS082222. This content is solely the responsibility of the authors and does not necessarily represent the official views of the National Institutes of Health.

Research reported in this publication was supported by National Institute of Drug Abuse National Institutes of Health under award number R44DA050357 This content is solely the responsibility of the authors and does not necessarily represent the official views of the National Institutes of Health.

### Appendix 1. Initial rate derivation

The initial rate of a biological process is the rate before it becomes limited by rate-limiting mechanisms. It is the rate of the process as time approaches zero. The parameters of the time course equation defining the initial rate can be identified by taking the limit of the equation as time approaches zero.

#### 1.1. Initial rate of model-free equations

The limit of the model-free equations (excluding Baseline) as time approaches zero is as follows:

Straight line, from equation (1):

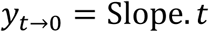

Association exponential, from equation (2):

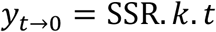

Rise-and-fall to baseline, from equation (3):

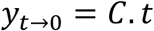

Rise-and-fall to steady-state, from equation (4):

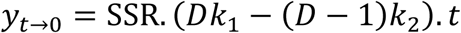

These are all straight line equations. The initial rate (IR) is the gradient of the line:

Straight line:

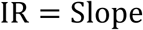

Association exponential:

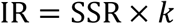

Rise-and-fall to baseline:

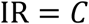

Rise-and-fall to steady-state:

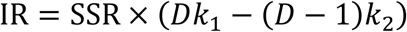

#### 1.2. Initial rate of kinetic model equations

Efficacy of the receptor-ligand complex is determined using a saturating concentration of agonist. The equations for the kinetic mechanistic models at saturating [*A*] a re g iven i n Supplementary Tables S1-S4. Taking the limit of these equations as time approaches zero, in all cases the equations reduce to the following:

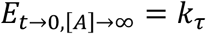

(Note in the case of equation (14) the limit is *k*_*τ*1_, the *k*_*τ*_ value of the non-desensitized receptor.) This finding indicates that the initial rate at saturating [*A*] from t he m odel-free a nalysis i s equivalent to *k*_*τ*_.

### Appendix 2. More complex models

Kinetic mechanistic models were developed here that incorporate receptor desensitization. These models employ the same formulation used in the kinetic mechanistic model ^32^. Agonist-bound receptor (*RA*) converts response precursor (*E*_*P*_) to the response (*E*) governed by the response generation rate constant *k*_*E*_. Here receptor desensitization is incorporated as a decrease of the receptor concentration that can generate the response; active receptor (*RA*) is converted to inactive receptor (*R*_0_*A*) governed by the desensitization rate constant *k*_*DES*_. Four models are considered – desensitization alone (Appendix 2.1); desensitization with response decay (Appendix 2.2); desensitization with resensitization and response decay (Appendix 2.3.1); and desensitization in which the desensitized receptor signals at a different rate (Appendix 2.3.2). Note the models assume the level of receptor-ligand complex does not change over time. This scenario likely applies to maximally-stimulating concentrations of ligand (used to quantify efficacy), and for all concentrations of lower potency ligands, as described in the Discussion. The model framework allows for extension to incorporate receptor-ligand binding kinetics as necessary ^32^.

#### 2.1. Desensitization alone model

The basic model is described by Scheme 1:

**Scheme 1.**
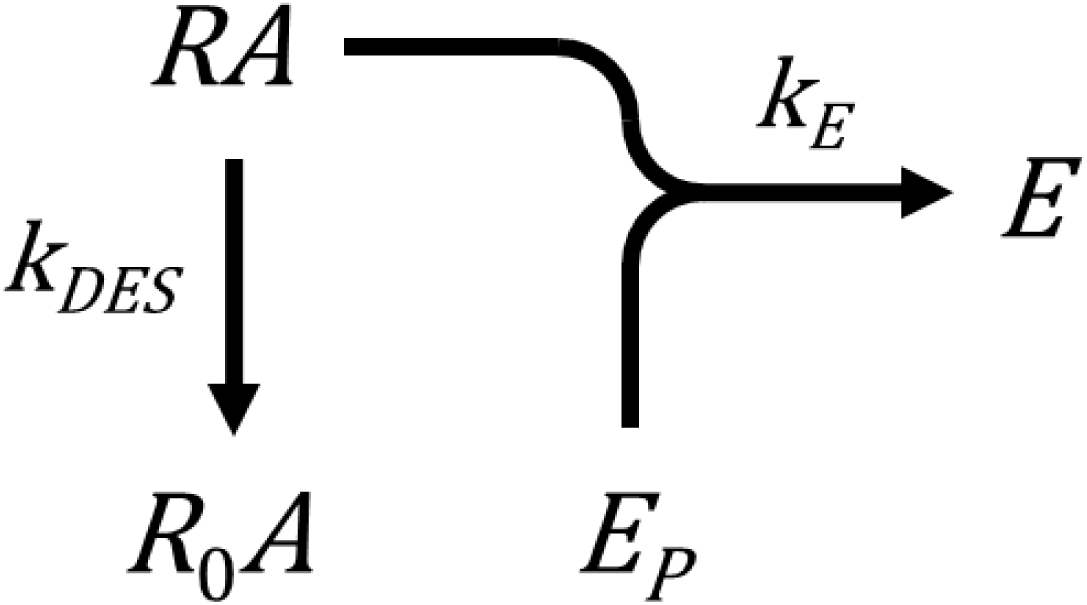

In this particular model, the response does not decay and the response precursor concentration remains constant over time, i.e. it is not depleted by generation of the response. The model is formulated for a maximally-stimulating concentration of agonist rather than a range of concentrations of agonist. This avoids the complexity of considering differential affinity for active versus desensitized receptors. The goal is an analytical equation for *E*, termed here an *E vs t* equation, which can be obtained using the Laplace transform method as follows. The differential equation for *E* is,

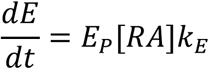

Since the concentration of agonist is saturating, [*RA*] does not change over time and is equal to the total concentration of non-desensitized receptors, termed [*R* _*a*_]_*TOT*_. The Laplace transform is then,

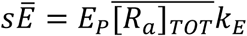

The differential equation for [*R*_*a*_] _*TOT*_ is,

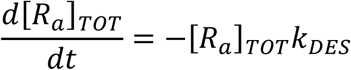

with boundary condition [*R*_*a*_]_*TOT,t*=0_ = [*R*]_*TOT*_. The Laplace transform for [*RA*] is then equation (10),

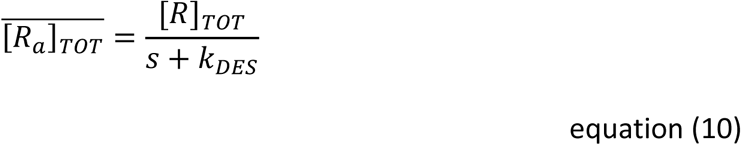

This is now substituted into the transform for *E* which gives, after re-arranging,

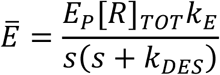

The *k*_*τ*_ term is now introduced. This is the initial rate of signal generation by the agonist-occupied receptor, the product of the total precursor concentration, total receptor concentration and the response generation rate constant ^32^, specifically:

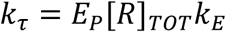

Substituting gives,

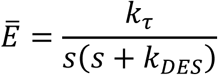

The analytic equation, equation (11) is now obtained by taking the inverse Laplace transform:

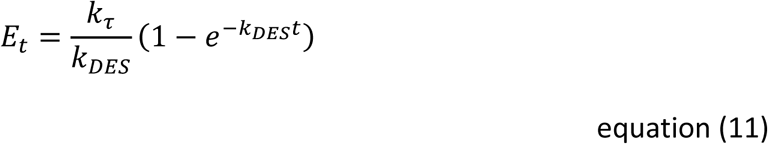

#### 2.2. Receptor desensitization with response degradation model

This model combines receptor desensitization with a second regulation of signaling mechanism, degradation of the response (for example, breakdown of second messenger molecules). (The response degradation model was described in the original kinetic mechanistic model ^32^.) The mechanism is described by Scheme 2:

**Scheme 2.**
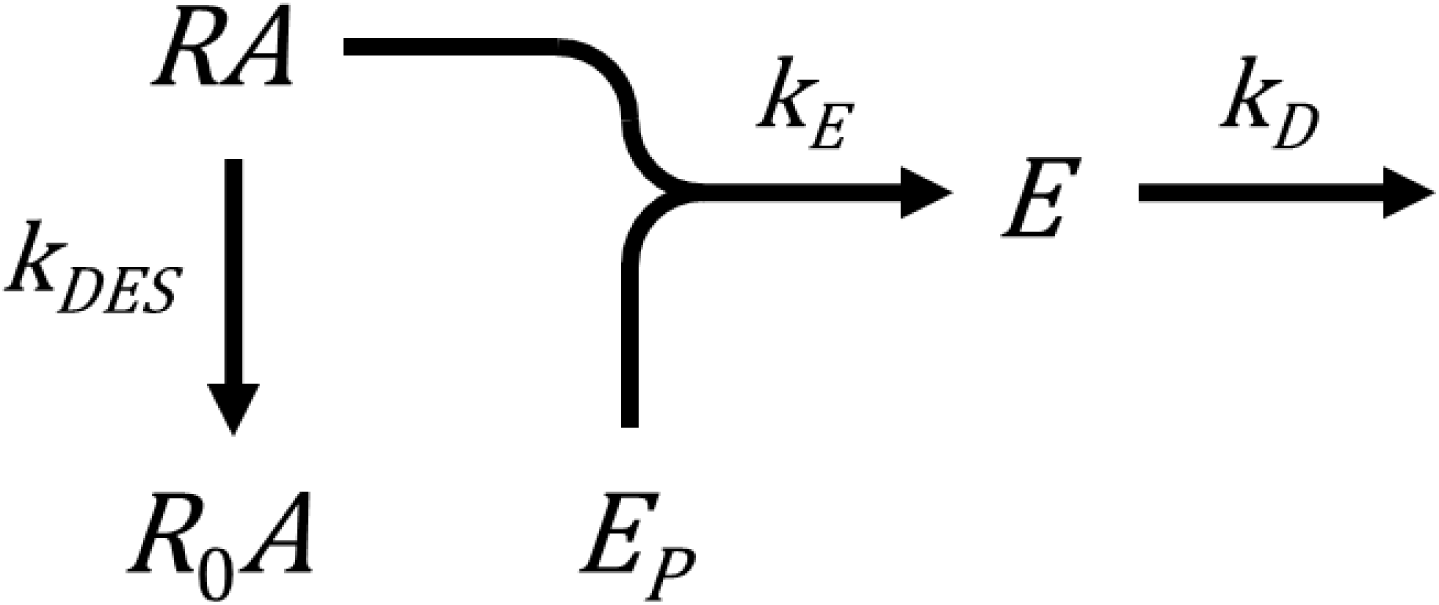

The *E vs t* equation is obtained, for a saturating agonist concentration, as follows. The differential equation and Laplace transform for *E* are,

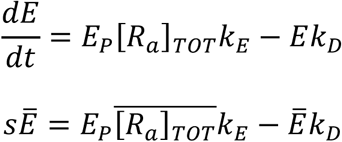

The derivation proceeds by taking the Laplace transform for [*R*_*a*_]_*TOT*_ (equation (10), Appendix 2.1) and substituting it into the transform for *E*, giving, after solving for *Ē*,

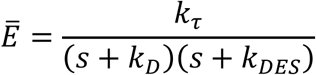

The *E vs t* equation is now obtained by taking the inverse Laplace transform, giving equation (12):

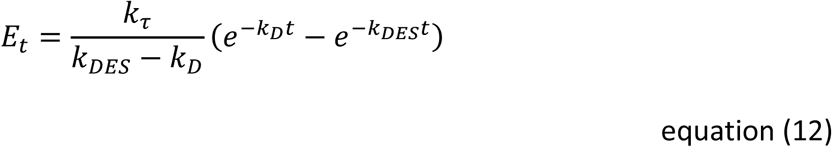

#### 2.3. Rise-and-fall to steady-state models

More complex regulation mechanisms have been described for GPCRs beyond the canonical receptor desensensitization and response degradation mechanisms. Three of these models are described and formulated here. All three reduce to a common general equation, the rise-and-fall to steady-state equation (equation (4)).

##### 2.3.1. Receptor desensitization and resensitization with response degradation

In this model, the receptor resensitizes after desensitizing. This is represented as a return to the active receptor state, *RA*, from the desensitized state *R*_0_*A*. This process proceeds at a rate defined by *k*_*RES*_, the resensitization rate constant. The mechanism, including response degradation, is an extension of the model in Appendix 2.2, and is represented by Scheme 3 below:

**Scheme 3.**
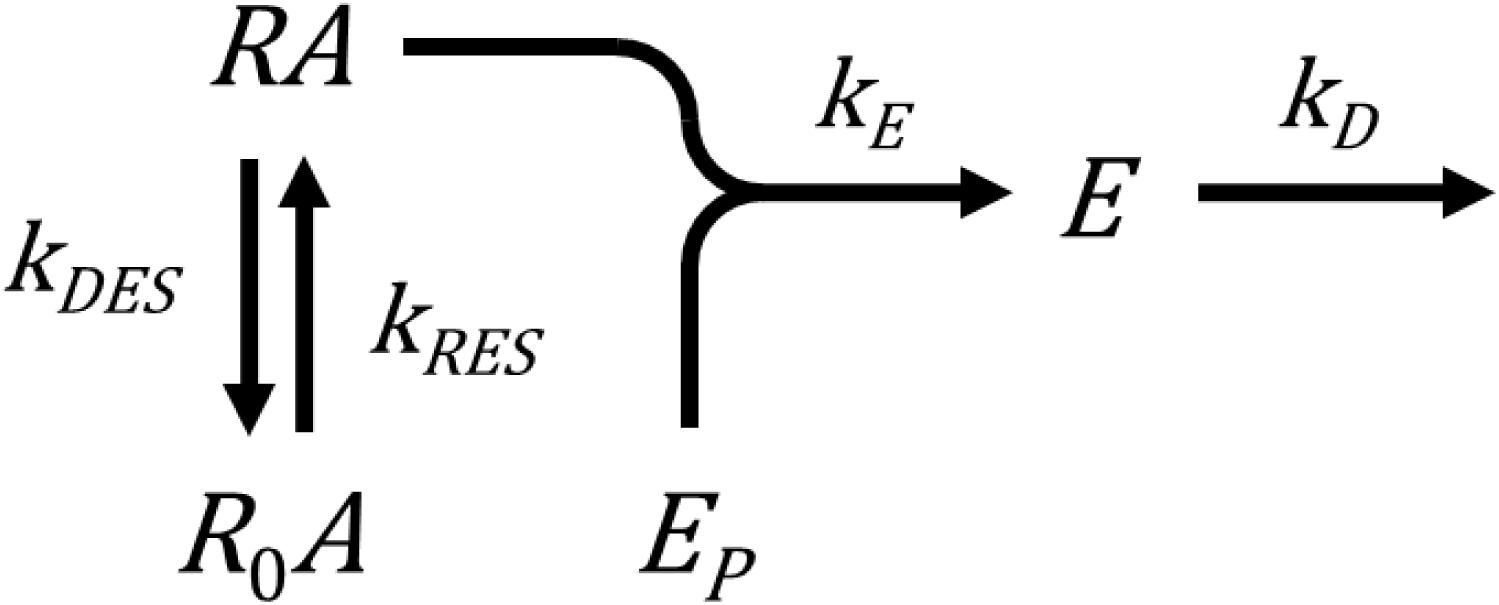

Here the model is formulated for a saturating concentration of agonist. This simplifies the model by avoiding consideration of the fate of desensitized receptors that become unbound by the agonist. The *E vs t* equation can be obtained using Laplace transforms as follows. First, the differential equation and Laplace transform for *E* are,

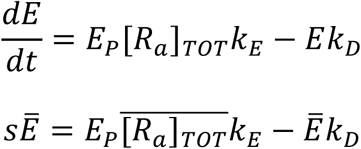

Note since we are dealing with a saturating concentration of agonist, [*RA*] is equal to [*R*_*a*_]_*TOT*_, the total concentration of non-desensitized receptors. The derivation proceeds by taking the Laplace transform for [*R*_*a*_]_*TOT*_ and substituting it into the transform for *E*. [*R*_*a*_]_*TOT*_ changes over time due to *RA* desensitization to *R*_0_*A* and resensitization of *R*_0_*A* back t o *RA*. Th e resu ltin g differential equation for [*R*_*a*_]_*TOT*_ is, for a saturating concentration of agonist,

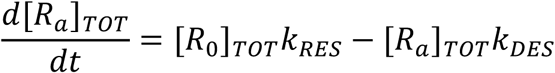

The [*R*_0_]_*TOT*_ term, the total concentration of desensitized receptors, can be replaced using the conservation of mass equation for the receptor ([*R*]_*TOT*_ = [*R*_*a*_]_*TOT*_ + [*R*_0_]_*TOT*_):

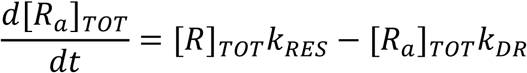

where *k*_*DR*_ = *k*_*DES*_ + *k*_*RES*_. The Laplace transform is,

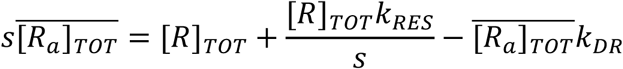

Note the transform includes [*R*_*a*_]_*TOT*_ at the initiation of the experiment and that this equals to the total concentration of receptors ([*R*]_*TOT*_) because desensitization has yet to take place. Solving for 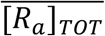 gives,

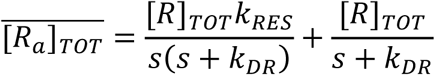

Substituting into the transform for *E* and solving for *Ē* gives,

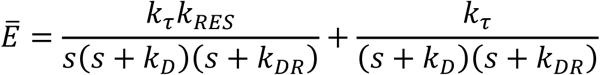

Taking the inverse Laplace transform gives the *E vs t* equation, equation (13):

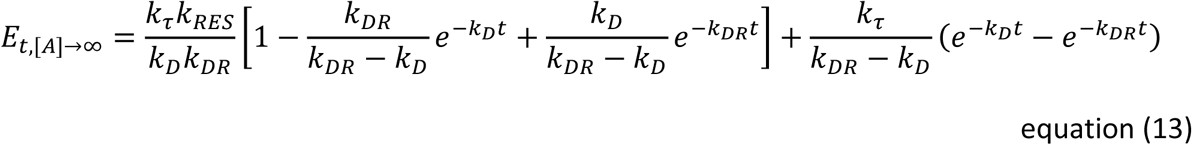

The equation can be rearranged to the general form:

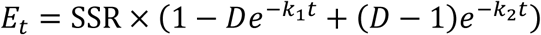

where SSR is the steady-state response, i.e. response as *t* → ∞. This rearrangement involves the intermediate step:

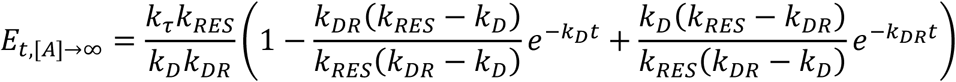

and the observation that the 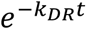 multiplier equals the 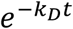 multiplier minus unity, i.e:

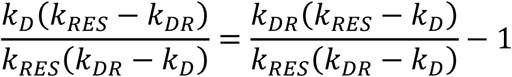

The parameters are defined as follows:

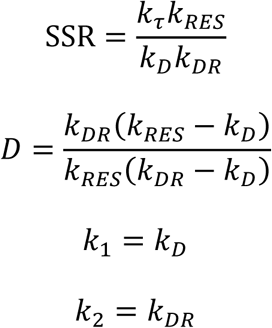

The initial rate of the response is *k*_*τ*_, shown by taking the limit of equation (13) as time approaches zero:

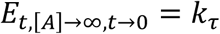

##### 2.3.2. Receptor desensitization, desensitized receptor signals

Recently it has been discovered that certain receptors which become desensitized can remain active for signaling. This mechanism is represented here by assuming the desensitized receptor *R*_0_*A* can couple to *E*_*P*_ to generate the response. The response generation rate constant of desensitized and non-desensitized receptor is assumed to be different (defined by *k*_*E*2_ and *k*_*E*1_, respectively). Here the simplest model is considered, one in which there is no receptor resesensitization, no precursor depletion, and a single rate of response degradation. It is represented by Scheme 4:

**Scheme 4.**
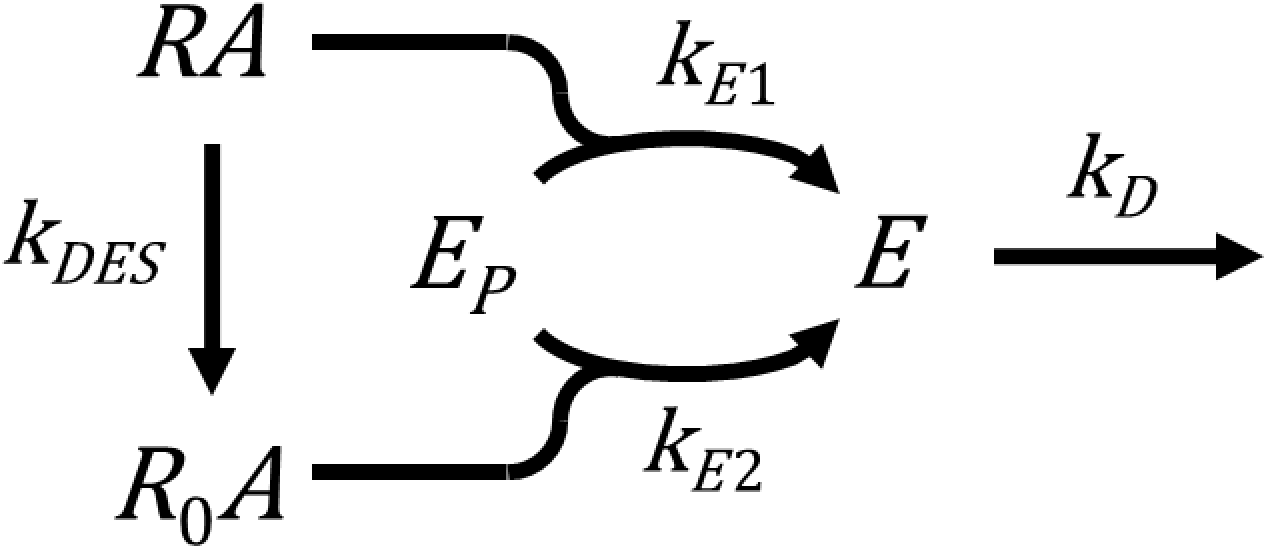

Here the model is formulated for a maximally-stimulating concentration of agonist. The differential equation for *E* is,

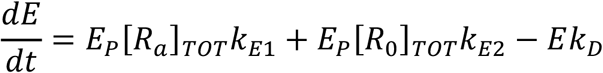

This can be simplified by employing the conservation of mass equation for the receptor, [*R*]_*TOT*_ = [*R*_*a*_]_*TOT*_ + [*R*_0_]_*TOT*_:

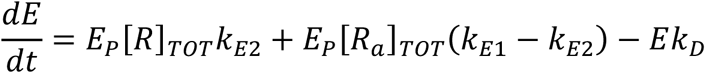

The Laplace transform is,

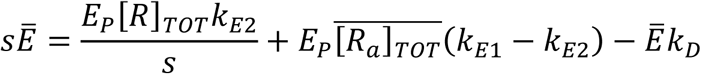

The Laplace transform for [*R*_*a*_]_*TOT*_ is equation (10) (Appendix 2.1):

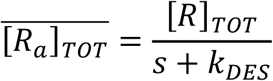

Substituting into the transform for *E* gives,

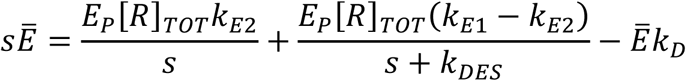

*k*_*τ*_ is now introduced. There are two terms, for non-desensitized and desensitized receptors, defined respectively as,

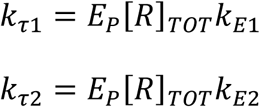

Substituting and solving for *Ē* gives,

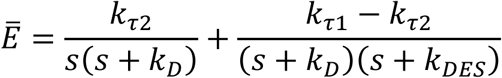

Taking the inverse Laplace transform gives the *E vs t* equation, equation (14)

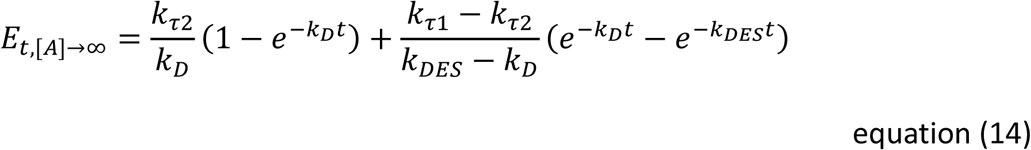

The equation can be rearranged to the general form:

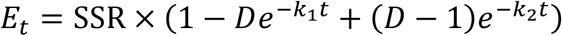

where SSR is the steady-state response, i.e. response as *t* → ∞. This rearrangement involves the intermediate step:

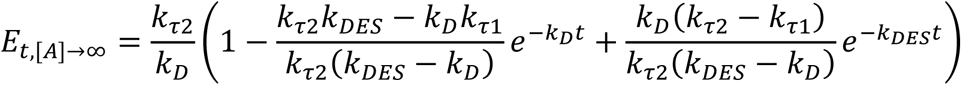

and the observation that the 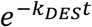 multiplier equals the 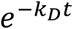 multiplier minus unity, i.e:

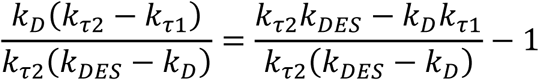

The parameters are defined as follows:

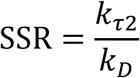

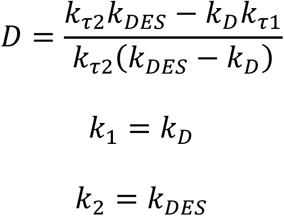

The initial rate of the response is *k*_*τ*1_, shown by taking the limit of equation (14) as time approaches zero:

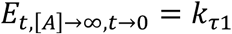

##### 2.3.3. Precursor depletion & response degradation to steady-state

In the calcium signaling mechanism, after the calcium rise and fall, the processes modulating cytoplasmic Ca^2+^ can reach a steady-state that results in a constant level of Ca^2+^ over time ^63,67-69^. The steady-state between Ca^2+^ mobilization and clearance can be accommodated within the model as an extension of the original precursor depletion and response degradation model (Model 4 in ^32^). The response degradation step becomes reversible. In other words, the response degradation product converts back to the response. This is represented by Scheme 5 below:

**Scheme 5.**
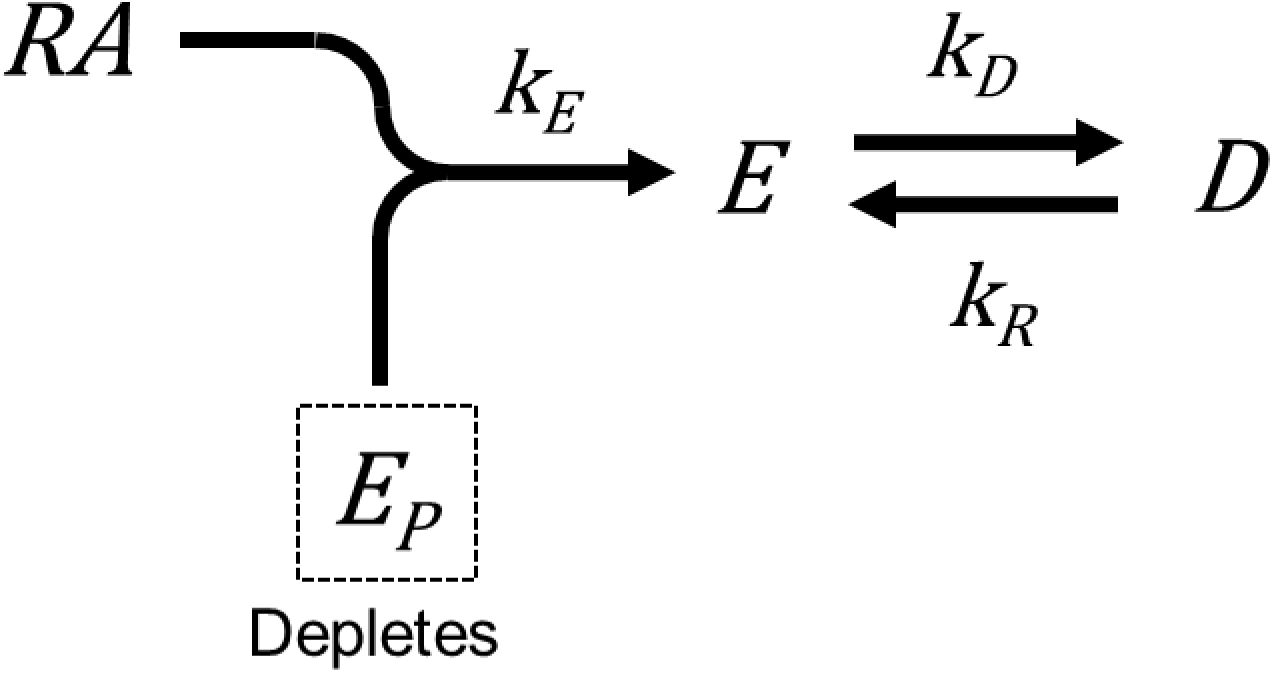

where *D* is the response degradation product and *k*_*R*_ is the response reformation rate constant. The differential equations for *E* is:

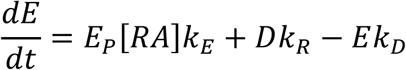

[*RA*] can be expressed as a function of the total receptor concentration:

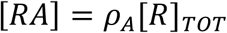

where *ρ*_*A*_ is fractional occupancy of receptor by agonist, given by the standard equilibrium receptor-ligand occupancy function:

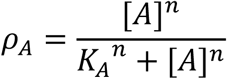

[*A*] is agonist concentration, *K*_*A*_ is the agonist-receptor equilibrium dissociation constant and *n* the slope factor.

*E*_*P*_ can be substituted using the conservation of mass equation, *E*_*P*(*TOT*)_ = *E*_*P*_ − *E* − *D*:

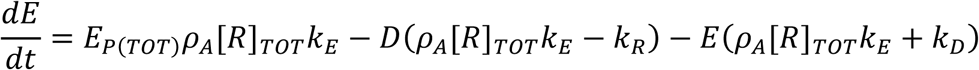

*D* can now be replaced with an expression in terms of *E* using Laplace transforms. The transform for *E* is,

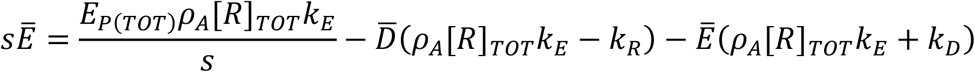

The differential equation and transform for *D* is,

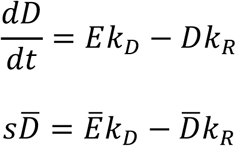

Solving for 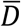 and substituting into the transform for *E* and solving for *Ē*, gives:

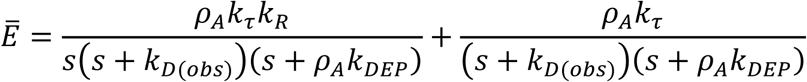

*k*_*DEP*_ is response precursor depletion rate constant, defined as the product of the total receptor concentration and the response generation rate constant, i.e. *k*_*DEP*_ = [*R*]_*TOT*_*k*_*E*_. The term *k*_*D*(*obs*)_ is *k*_*R*_ + *k*_*D*_. Taking the inverse Laplace transform gives the *E vs t* equation, equation (15):

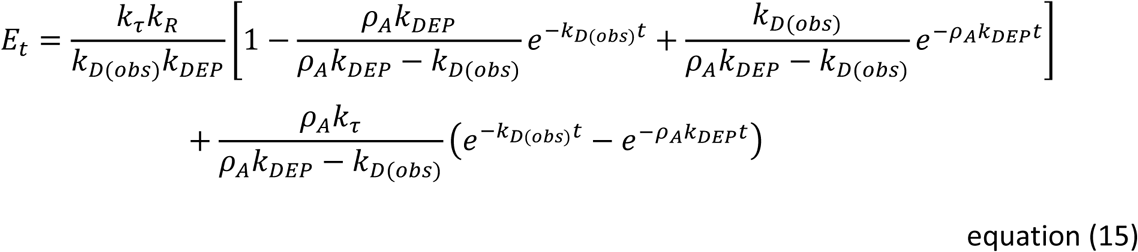

The equation can be rearranged to a general form:

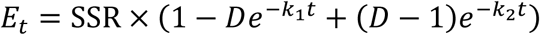

where SSR is the steady-state response, i.e. response as *t* → ∞. This rearrangement involves the intermediate step:

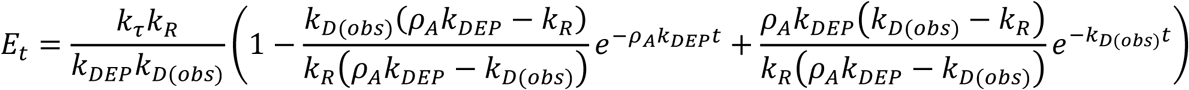

and the observation that the 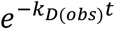 multiplier equals the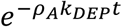 multiplier minus unity, i.e:

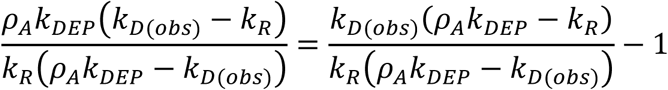

The parameters are defined as follows:

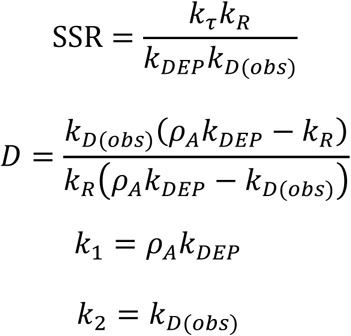

At a saturating concentration of agonist, parameters that are agonist dependent (*D* and *k*_1_) are defined as follows:

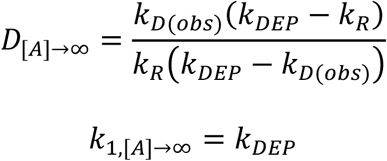

The expanded equation at a saturating concentration of agonist is equation (16):

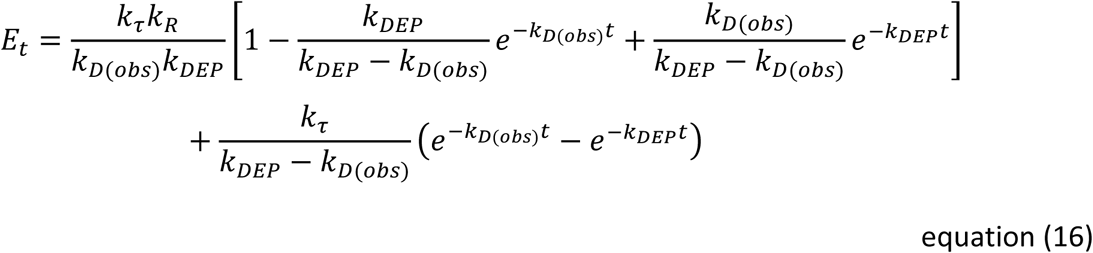

The initial rate of the response is *k*_*τ*_, shown by taking the limit of equation (16) as ti me approaches zero:

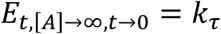

## Methods

### Extraction of time course data

Data were extracted from figures in published articles using a plot digitizer (Graph Grabber v2, Quintessa Limited, Henley-on-Thames, UK).

### Equations for model-free analysis

For model-free analysis, time course data were fit to empirical time course equations. The following equations were used when the response started at the moment of ligand addition. The straight line equation is equation (1),

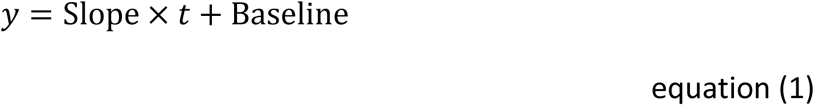

where *y* is response, Slope is the gradient of the line and *t* is the time of response measurement. Baseline is the response in the absence of ligand. Note Baseline is assumed to remain constant over time. The corresponding equation in Prism is named “Straight line” and the parameter Yintercept in this equation corresponds to Baseline ^90^.

The association exponential equation is equation (2):

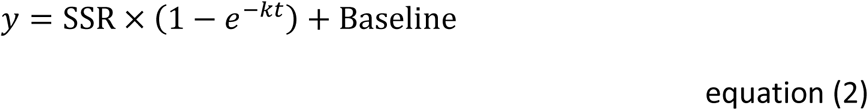

where SSR is the steady-state response, specifically the ligand-specific response as time approaches infinity. Note the *y* asymptote value as time approaches infinity is SSR + Baseline. *k* is the observed rate constant in units of *t*^-1^. The corresponding equation in Prism 8 is named “One-phase association,” ^91^ in which Span corresponds to SSR, Y0 corresponds to Baseline, and Plateau corresponds to SSR + Baseline.

The rise-and-fall to baseline equation is equation (3):

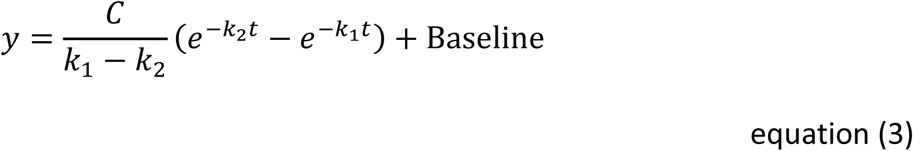

where *C* is a fitting constant in units of y-units.*t*^-1^ (which is the initial rate of signaling, see Appendix 1.1) and *k*_1_ and *k*_2_ are observed rate constants in units of *t*^-1^. In the analysis, *k*_1_ is the larger of the two rate constant values, constrained to be greater than *k*_2_ (see “Fitting procedures” below). This equation has been loaded into a Prism template as a User-defined equation named “Rise-and-fall to baseline time course” (see “Signaling kinetic model-free equations” file available in the supplementary files).

The rise-and-fall to steady-state equation is equation (4):

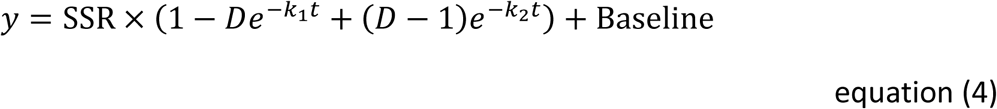

where, as above, SSR is the steady-state response, specifically the ligand-specific response as time approaches infinity. Note the *y* asymptote value as time approaches infinity is SSR + Baseline. *D* is a unitless fitting constant, and *k*_1_ and *k*_2_ are observed rate constants in units of *t*^-1^. In the analysis, *k*_1_ is the larger of the two rate constant values, constrained to be greater than *k*_2_ (see “Fitting procedures” below). This equation has been loaded into a Prism template as a user-defined equation named “Rise-and-fall to steady state time course” in which the parameter SteadyState corresponds to SSR (see “Signaling kinetic model-free equations” file available in the supplementary files).

In certain circumstances it is desirable to float the initiation of signaling time in the analysis. This enables the analysis to accommodate slight uncertainty as to the precise time point of ligand addition, especially for rapidly-generated signals. It also allows for signaling mechanisms where there is a delay between ligand-receptor binding and initiation of detectable signaling, often observed in gene expression assays, and thought to be due to a necessary build-up of signal transduction intermediates to a threshold level ^47^. When floating the start time in the analysis it is necessary to establish the baseline response level prior to the start time. Equations 1-4 above can be adapted to incorporate a fitted start time, the corresponding equations being equations (5-8) below:

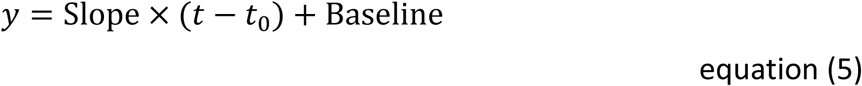

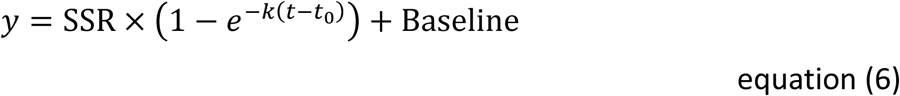

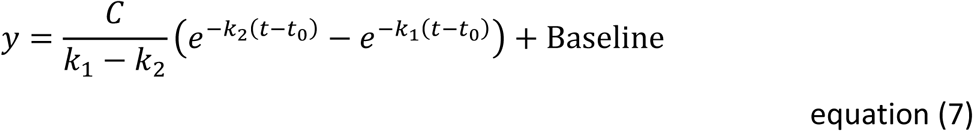

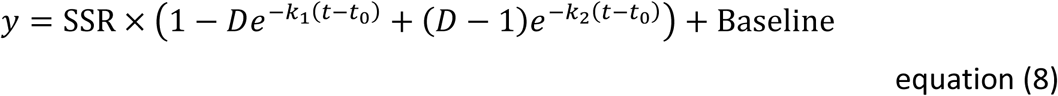

where *t*_0_ is the signal initiation time and *t* is time uncorrected for the signal initiation time. Equation (6) is equivalent to the equation named “Plateau followed by one phase association” in Prism 8, in which Span corresponds to SSR and Y0 corresponds to Baseline ^92^. Equations 5,7 and 8 have been loaded into a Prism template as User-defined equations named “Baseline then straight line time course”, “Baseline then rise-and-fall to baseline time course” and “Baseline then rise-and-fall to steady state time course” respectively (see “Signaling kinetic model-free equations” Prism file available in the supplementary files). In Prism the equations are coded with an IF statement to fit the baseline before *t*_0_. For example, equation (7) is written in Prism as,

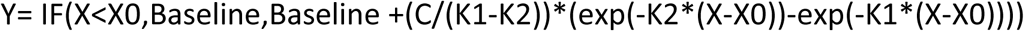

In some cases, stimulation of signaling results in a decrease of response signal. In other words, the detected response signal decreases over time. This can result from technical considerations, for example, certain biosensors decrease in signal upon binding the response analyte (so called “Downward” sensors). It can also result from the signaling mechanism, for example stimulation of G_i_ is often detected by the resulting inhibition of cAMP accumulation. This can be handled by taking the inverse of the detected signal and plotting this versus time (creating an “Upward” time course, see for example Supplementary Fig. S2d). Alternatively, downward data can be fit to the downward analogue of the equations. Examples of this approach are used in this study (inhibition of cAMP accumulation, see Supplementary Fig. S2f;, and GIRK channel currents, see Supplementary Fig. S2h); data were analyzed with the downward analogue of the association exponential equation, which is the exponential decay equation shown below (equation (9)):

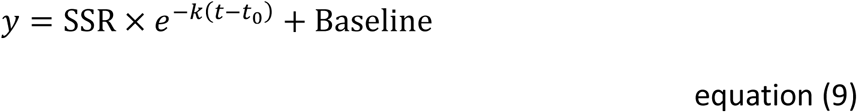

Note the response at *t* = 0 is equal to SSR + Baseline. The corresponding equation in Prism 8 is named “Plateau followed by one phase decay,” in which Span corresponds to SSR, Plateau corresponds to Baseline, and Y0 corresponds to SSR + Baseline ^93^.

### Calculation of initial rate

The initial rate of signaling was determined by first fitting the data to the time course equations, then using the fitted parameter values to calculate the initial rate (IR). This calculation utilized an equation defining the initial rate in terms of the fitted parameters, the equation obtained by taking the limit of the time course equation as time approaches zero (the formal definition of the initial rate condition). These equations were derived in Appendix 1:

Straight line:

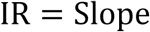

Association exponential:

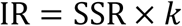

Rise-and-fall to baseline:

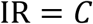

Rise-and-fall to steady-state:

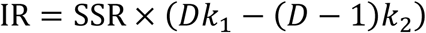

### Equations for kinetic mechanistic model analysis and simulation

The kinetic mechanistic model was used to simulate and analyze time course data using the model equations, derived in ref. ^32^ and Appendix 2. The equations employed are given in Supplementary Tables S1-S4 and are listed in Supplementary information. A Prism template containing the equations called “Signaling kinetic mechanism equations” is available in the supplementary files. The terms in the models are defined in Supplementary Table S5. Data in Fig. 5a (Ca^2+^ mobilization via the GnRH_1_ receptor) were analyzed using the equation for the precursor depletion & response degradation to steady-state model (Appendix 2.3.3) that incorporated a variable signal initiation time (*t*_0_):

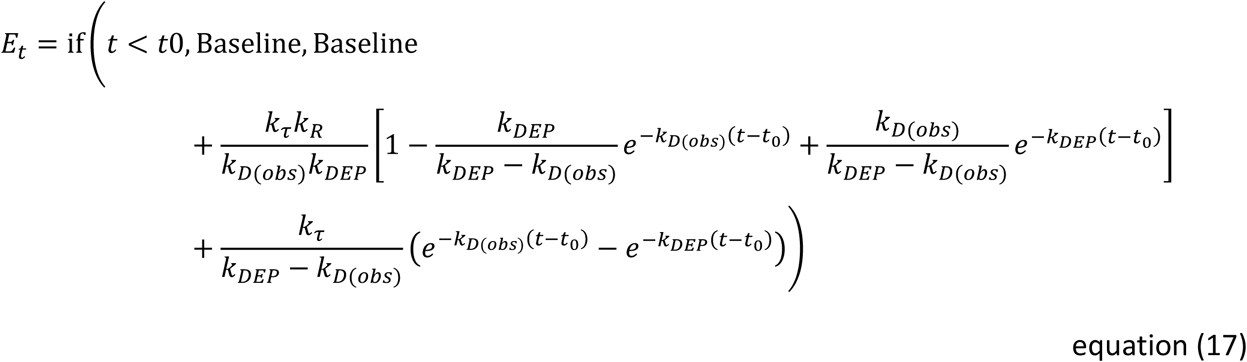

where,

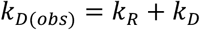

### Fitting procedures

Data were fit by nonlinear regression to the equations using Prism 8. In almost all cases, the default fitting settings were used, specifically least-squares nonlinear regression with medium convergence criteria, no special handling of outliers and no weighting ^72^. In two cases, Fig. 4b and Fig. 5b, the default settings yielded an ambiguous fit, in that some of the parameter values were not resolved (see “Curve fit results” Excel file in Supporting Material) ^73^. In these cases, unambiguous parameter values were obtained using the “Robust regression” option which minimizes the contribution of outliers to the fit ^74^. In the rise-and-fall fits (equations 3,4,7 and 8), which contain two rate constants *k*_1_ and *k*_2_, *k*_1_ was assumed to be the larger of the rate constant values and this was handled by constraining *k*_1_ to be greater than *k*_2_. In all cases, rate constant values were constrained to be greater than zero. The fitted values, the standard error and the correlation coefficient R^2^ are given in the “Curve fit results” Excel file in Supporting Material. Values were given to four significant figures. When the signal initiation time was floated (equations 5-9), the initial value (X0) was entered manually.

In order to obtain an estimate of the error of the fitted parameters, the standard error was computed as described ^94^, assuming a symmetrical confidence interval. This required changing the default “Calculate CI or parameters” from asymmetrical to symmetrical in the “Confidence” dialogue ^95^.

The baseline response, that in the absence of ligand, was handled by incorporating a baseline term into the equations and fitting this parameter as part of the analysis. A commonly-used alternative, subtracting baseline from the data set and excluding the baseline term from the equation, was found to distort the fit significantly for the rise-and-fall to baseline equation, unless the subtracted baseline value was in very close agreement with that determined by incorporating the baseline into the curve-fitting procedure.

For the V_2_ vasopressin receptor experiments, the biosensor fluorescence intensity data were normalized to baseline. Specifically, response was quantified as the fluorescence after ligand addition divided by the mean baseline signal before addition, a metric termed F / ΔF. (The mean baseline signal is the mean of the fluorescence intensity measurements before ligand addition.) The data was handled in this way because it provides a convenient intra-well control, minimizing error due to any slight well-to-well differences in the amount of sensor or number of cells. The arrestin sensor data was inverted by calculating the value 1 - F / ΔF. This was done because the arrestin sensor is a downward sensor (a decrease in fluorescence resulted from receptor interaction) ^26^. In the fitting procedure, each technical replicate was considered an individual point.

Concentration-response data for the initial rate were fit to a sigmoid curve equation, the “Log(agonist) vs. response -- Variable slope” equation in Prism ^96^:

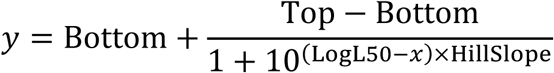

The “Bottom” parameter was constrained to zero. (Note in the Prism formulation, L_50_ is written as EC_50_. The EC_50_ term is not used here because it has an explicit pharmacological definition ^57^.)

### Measurement of cAMP generation and arrestin recruitment using biosensors

Genetically-encoded biosensors were used to measure cAMP generation and arrestin recruitment via the V_2_ vasopressin receptor. The sensors have been described previously (red cADDis for cAMP ^77^ and see ref ^26^. for arrestin recruitment), and are packaged in the BacMam vector for delivery to cells. The cDNA for the V_2_ vasopressin receptor was obtained from the cDNA Resource Center (Bloomsburg University, Bloomsburg, PA). The experiments were conducted in HEK293T cells transfected with the receptor and each sensor individually (i.e. one batch of cells with receptor and the cAMP sensor and a separate batch with receptor and arrestin sensor). HEK 293T cells were cultured in Eagle’s minimum essential media (EMEM) supplemented with 10% fetal bovine serum (FBS) and penicillin-streptomycin at 37 °C in 5 % CO_2_. One day before the transfection, HEK293T cells were seeded at a density of 27,000 cells/well in a 96-well plate (Greiner CELLCOAT® #655946). Approximately 16-20 hours later, cells were transfected with 50 ng of the V_2_ receptor plasmid per well, using Lipofectamine 2000 from Invitrogen (Waltham, MA USA). After an approximately 4 hourr incubation period at 37 °C and 5 % CO_2_, the transfection mix was replaced with 100uL complete cell culture media and the cells were allowed to recover for approximately 1.5 hours under normal growth conditions. The cells were then transduced with either the red caDDis or green arrestin sensor BacMam stocks. To prepare the transduction mixture, the BacMam containing cADDis or the arrestin sensor, 2 mM sodium butyrate (Sigma), and EMEM were combined in a final volume of 50 μL. For each experiment, 6.18 × 10^8^ viral genes of cADDis virus or 4.24 × 10^8^ viral genes of arrestin sensor virus were added to each well. The 50 uL transduction mixture was then added to the 96-well plate (50 uL/well) and incubated for approximately 24 hours at 37 °C in 5 % CO_2_.

Thirty minutes prior to fluorescence plate reader experiments, the media in each well was replaced with 150 μL of Dulbecco’s phosphate buffered saline (DPBS) supplemented with Ca^2+^ (0.9 mM) and Mg^2+^ (0.5 mM). Fluorescence plate reader experiments were performed on the Synergy Mx reader (BioTek, Winooski, VT). The green fluorescence detection was recorded using 488/20 nm excitation and 525/20 nm fluorescence emission, while red fluorescence detection was recorded using 565/20 nm excitation wavelength and 603/20 nm fluorescence emission. After acquiring the baseline fluorescence for several minutes, drug was added manually with a multichannel pipette in a volume of 50 µL at the indicated time points.

Oxytocin and vasopressin were obtained from Cayman Chemical. All working concentration of drugs were dissolved in DPBS and added manually to the HEK 293T cells at the indicated concentrations and time points.

## Author contributions statement

S.R.J.H. and L.J.B. designed the data analysis framework and analyzed data. P.H.T, A.M.Q. and T.E.H. conceived, designed and executed experiments and analyzed data. S.R.J.H wrote the manuscript. All authors reviewed the manuscript.

## Competing Interests statement

The authors declare no competing interests.

## Supplementary Information

### List of Supporting Material files

#### Curve fit results

Curve fit results details for the analyses in the figures

#### Kinetic mechanism model simulations

Excel 2016 file

Time course data simulator for the mechanistic kinetic models

#### Signaling kinetic model-free equations

Prism 8 file

Template containing model-free equations

#### Signaling kinetic mechanism equations

Prism 8 file

Template containing kinetic mechanism model equations

### Supplementary Figures

**Supplemental Figure S1.**
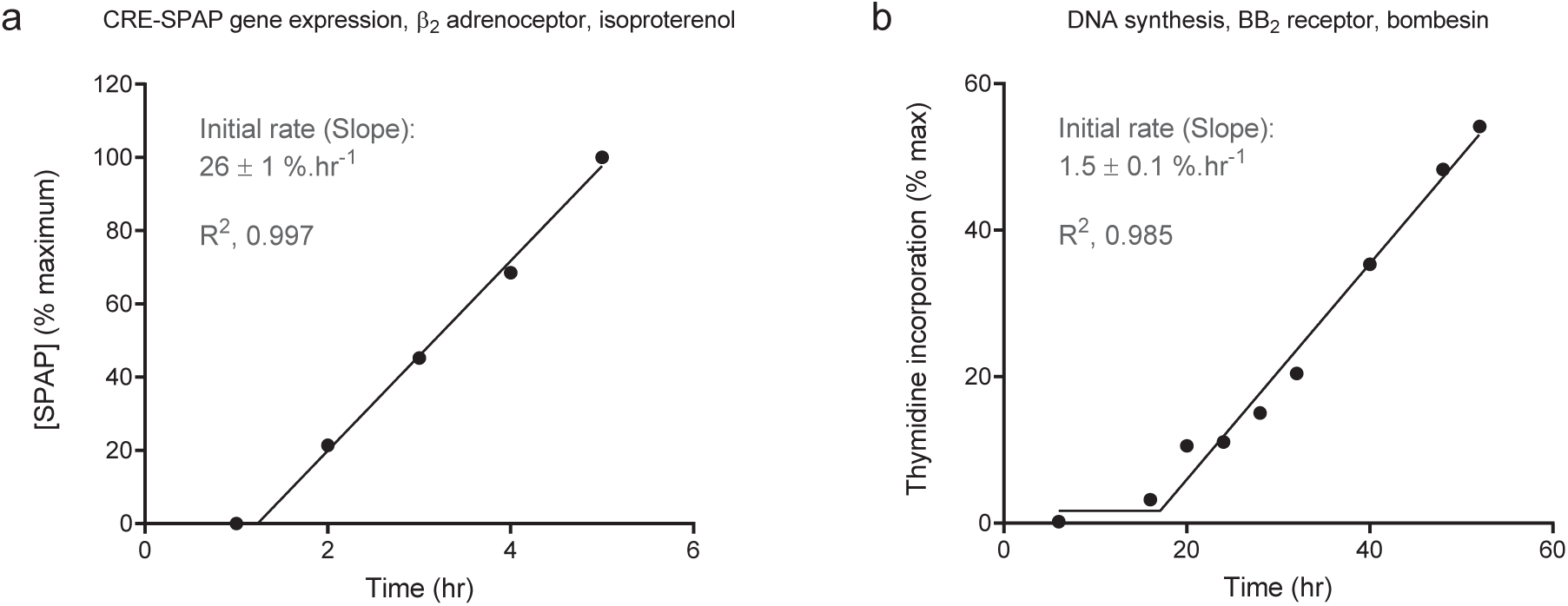
Additional linear time course profiles of GPCR signaling. (a) Gene expression induction (CRE-SPAP production) via the β_2_ adrenoceptor stimulated by 10 μM isoproterenol (data from Table 2 of ^1^). (b) DNA synthesis in Swiss 3T3 cells stimulated by 10 nM bombesin via the BB_2_ receptor (data from Fig. 2 of ^2^). Data were fit to equation (5), which incorporates a delay between application of ligand (at t = 0) and initiation of response. The Slope value is the fitted value ± the fit SEM ^3,4^.

**Supplemental Figure S2.**
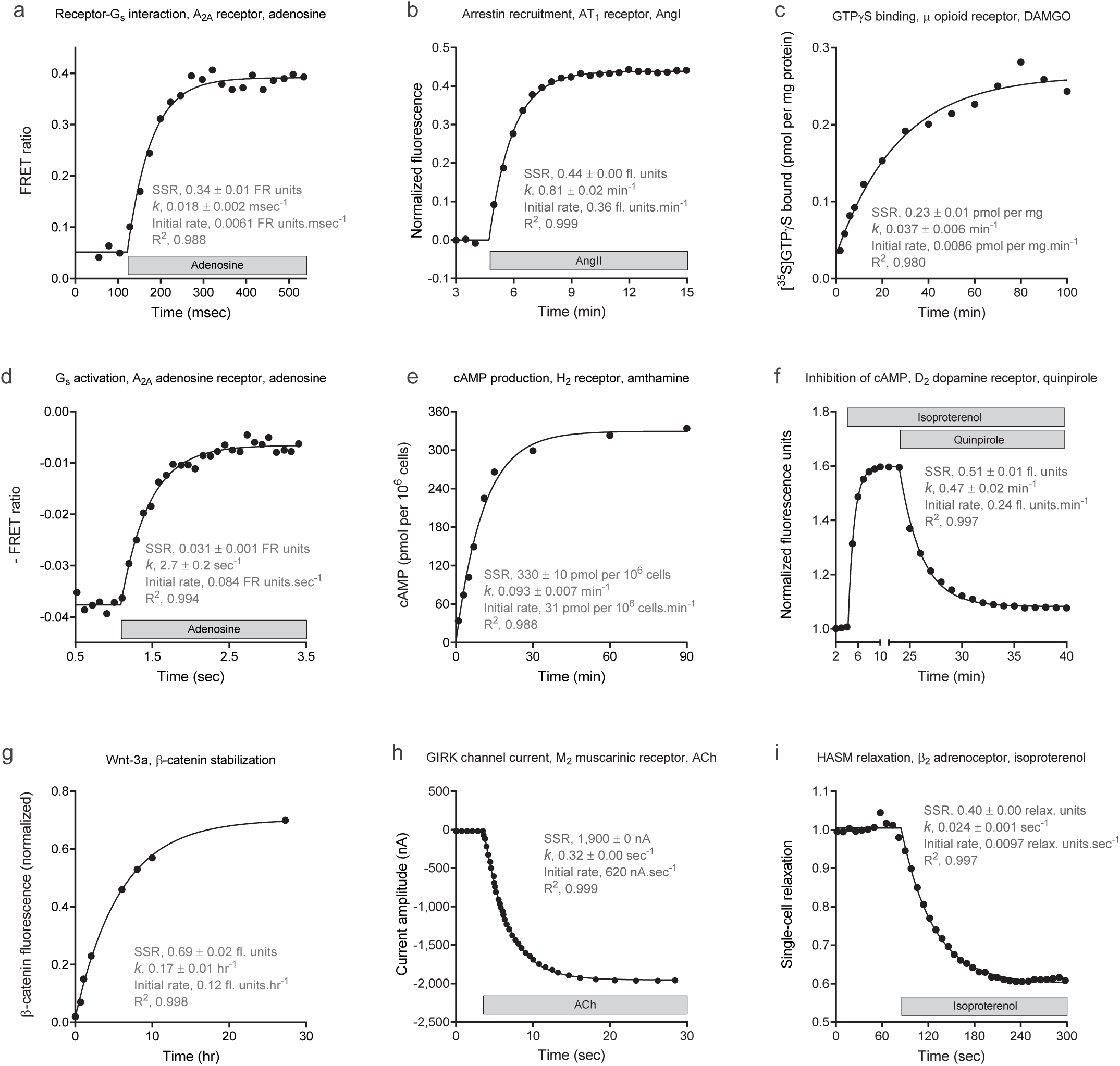
Additional association exponential time course profiles of GPCR signaling. (a) Gs interaction with the A_2A_ adenosine receptor stimulated by 1 mM adenosine, measured by FRET (data from Fig. 1c of ^5^, fit to equation (6)). (b) Arrestin recruitment to the AT_1_ angiotensin receptor stimulated by 32 μM AngII (data from Fig. 4a of ^6^, fit to equation (6)). (c) DAMGO-stimulated [^35^S]GTPγS binding via the μ opioid receptor in C6 cell membranes (1 μM DAMGO, data from Fig. 1 of ^7^, fit to equation (2)). (d) G_s_ activation by the A_2A_ receptor stimulated with 100 μM adenosine, measured by FRET (data from Fig. 2c of ^5^, note the inverse of the FRET ratio is used). (e) cAMP production via the H_2_ histamine receptor in U-937 cells stimulated by 10 μM amthamine (data from Fig. 4, D5 cell line, of ^8^, fit to equation (6), 1 mM isobutylmethyxanthine present). (f) Inhibition of cAMP production via the D2 dopamine receptor in HEK293T cells activated by 100 nM quinpirole. cAMP production was first stimulated by 1 nM isoproterenol via the endogenously-expressed β_2_ adrenoceptor. After the plateau had been reached, quinpirole was applied and inhibition recorded. [Data are from Fig. 4a of ^9^, fit to equation (9) (quinpirole plus isoproterenol phase) and equation (6) (isoproterenol alone phase). Note the *y* values are the inverse of those in the published figure. (g) β-catenin stabilization in L cell fibroblasts stimulated by 400 ng/ml Wnt-3a (data from Fig. 4d of ^10^, fit to equation (2)). (h) GIRK channel gating by 1 μM acetylcholine via the M_2_ muscarinic acetylcholine receptor (data from Fig. 1a of ^11^, fit to equation (6)). (i) Relaxation of human airway smooth muscle cells by 10 μM isoproterenol via the β_2_ adrenoceptor (data from Fig. 1a of ^12^, fit to equation (6)). The fitted values SSR (steady-state response) and *k* (the rate constant) are the fitted value ± the fit SEM ^3,4^. The initial rate was calculated as the SSR multiplied by *k*. The grey bar indicates the time interval of agonist application.

**Supplemental Figure S3.**
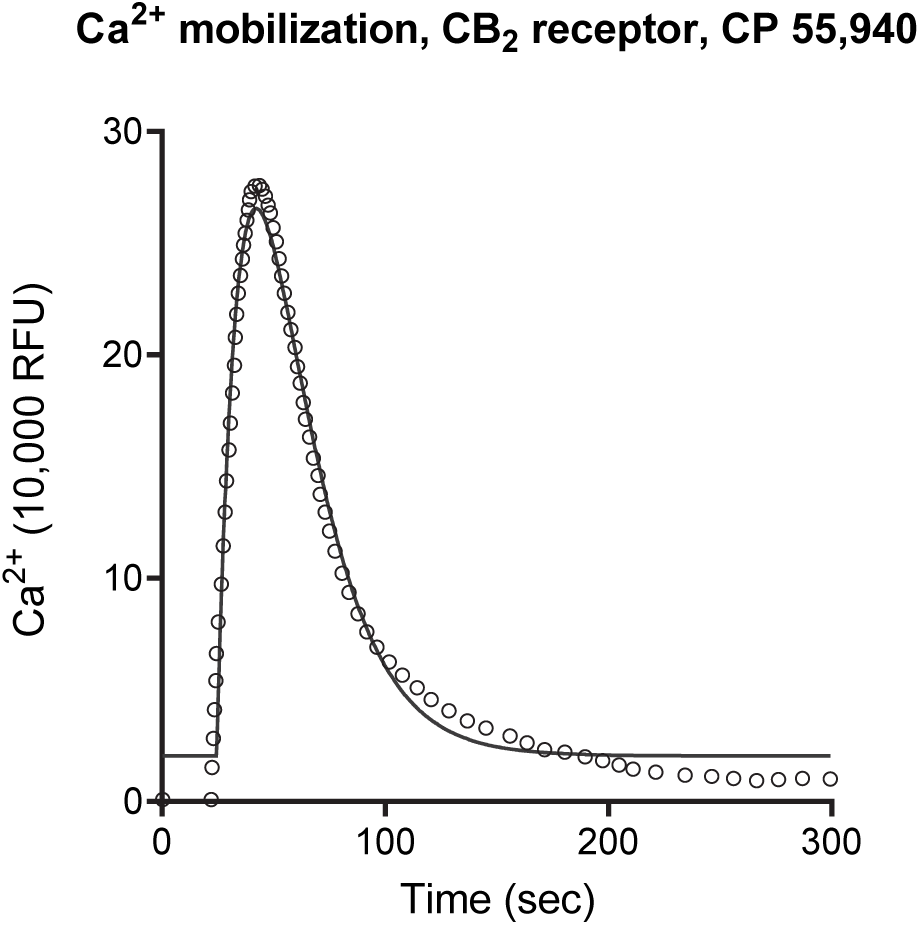
Example of rise-and-fall time course data that do not fit to the model-free equations used in this study: Ca^2+^ mobilization via the 5HT_7_ receptor stimulated by 10 μM CP 55,940. The curve is the fit to the rise-and-fall to baseline equation (equation (7)). Data are from Fig. 4a of ^13^.

**Supplemental Figure S4.**
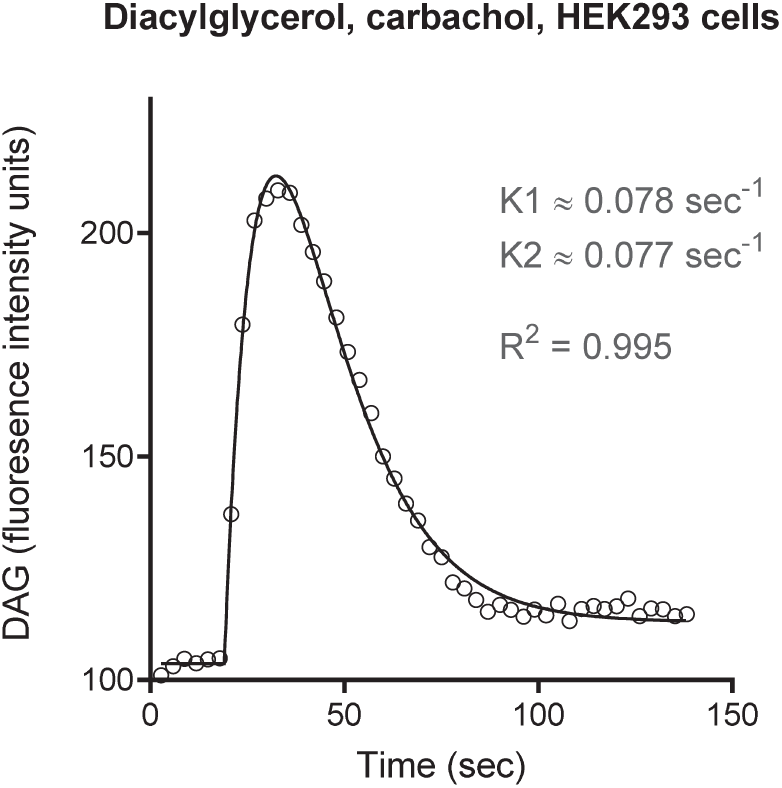
Example of rise-and-fall time course profile in which the two rate constant values are almost equal and convergence was not reached: Diacylglycerol production stimulated by carbachol in HEK293 cells (data from Fig. 2A (upper panel) of ^14^, fit to equation (8)). For fit details, see “Curve fit results” Excel file, “Rise-and-fall to steady-state” worksheet, in Supporting Material.

### Supplementary Tables

**Supplementary Table S1.**
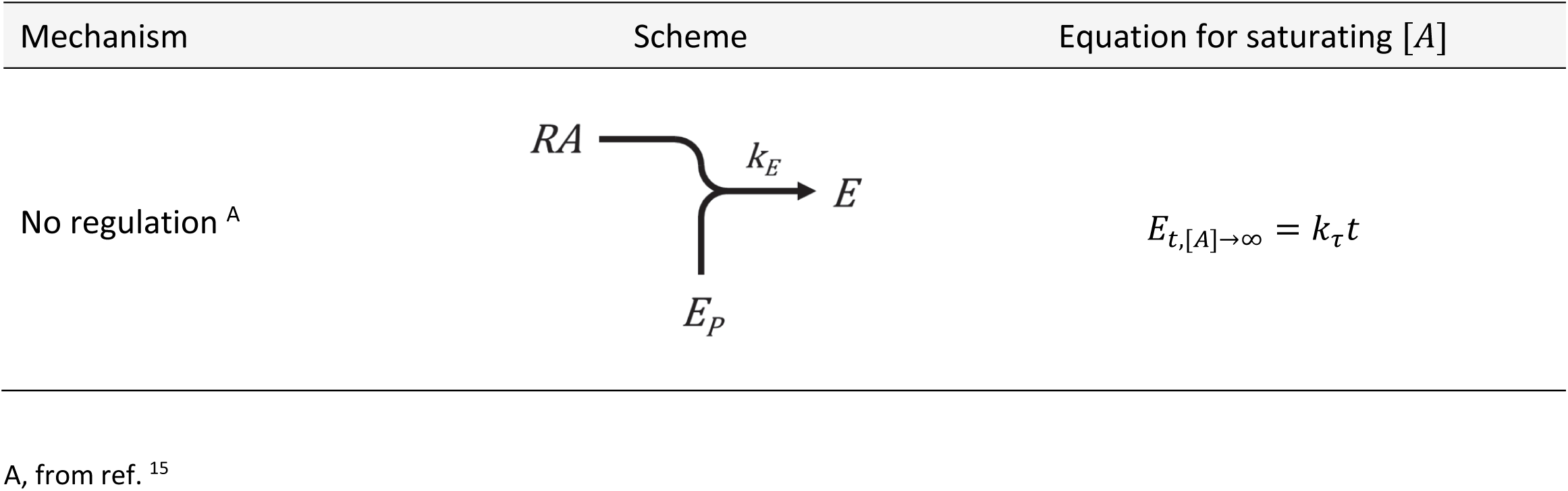
Summary of mechanism for straight line time course profile.

| Mechanism | Scheme | Equation for saturating [A] |
| --- | --- | --- |
| No regulation <sup>A</sup> | | $E_{t,[A] \rightarrow \infty} = k_{\tau} t$ |

A, from ref. <sup>15</sup>

**Supplementary Table S2.**
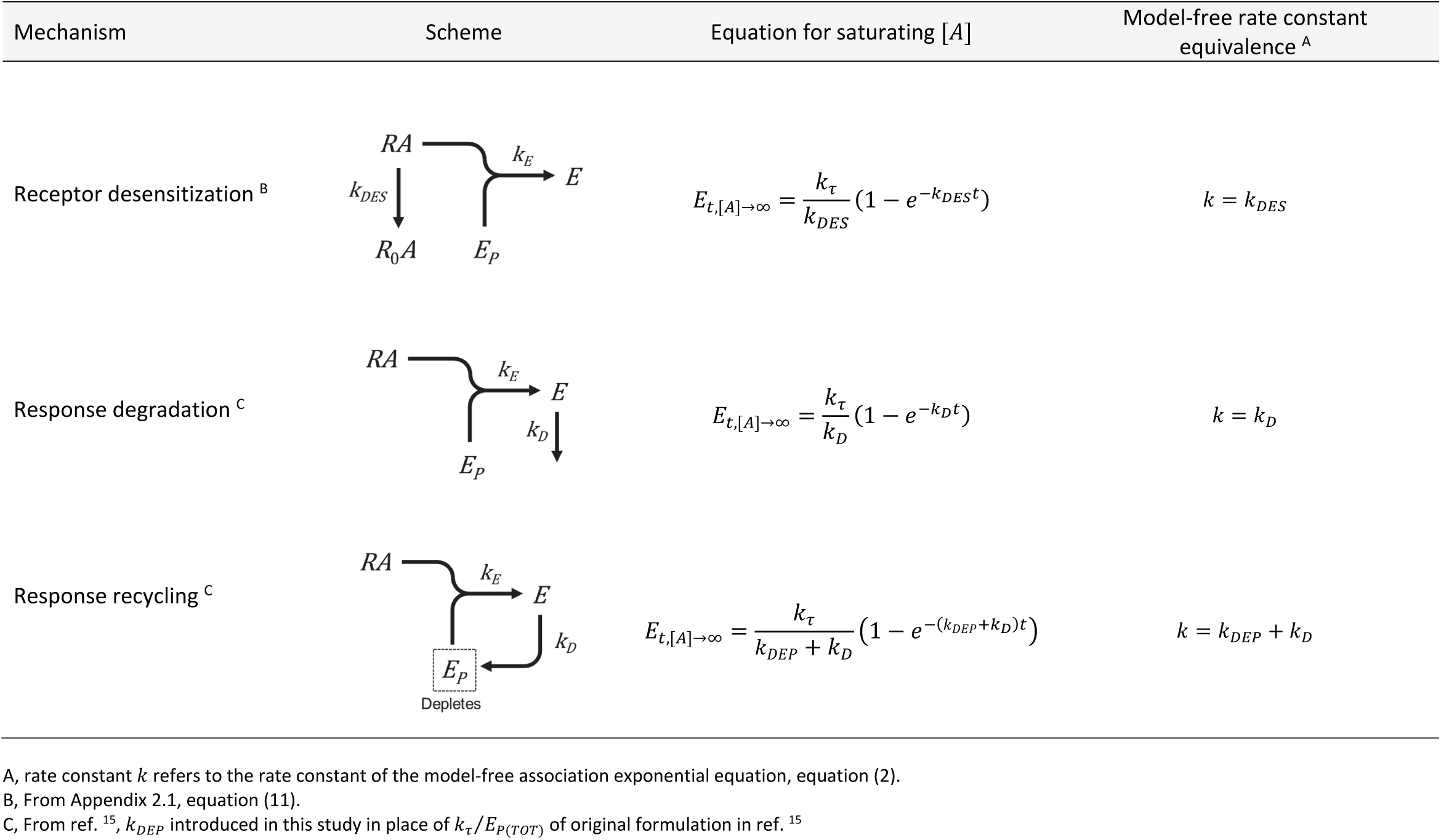
Summary of mechanisms for association exponential time course profile.

**Supplemental Table S3.**
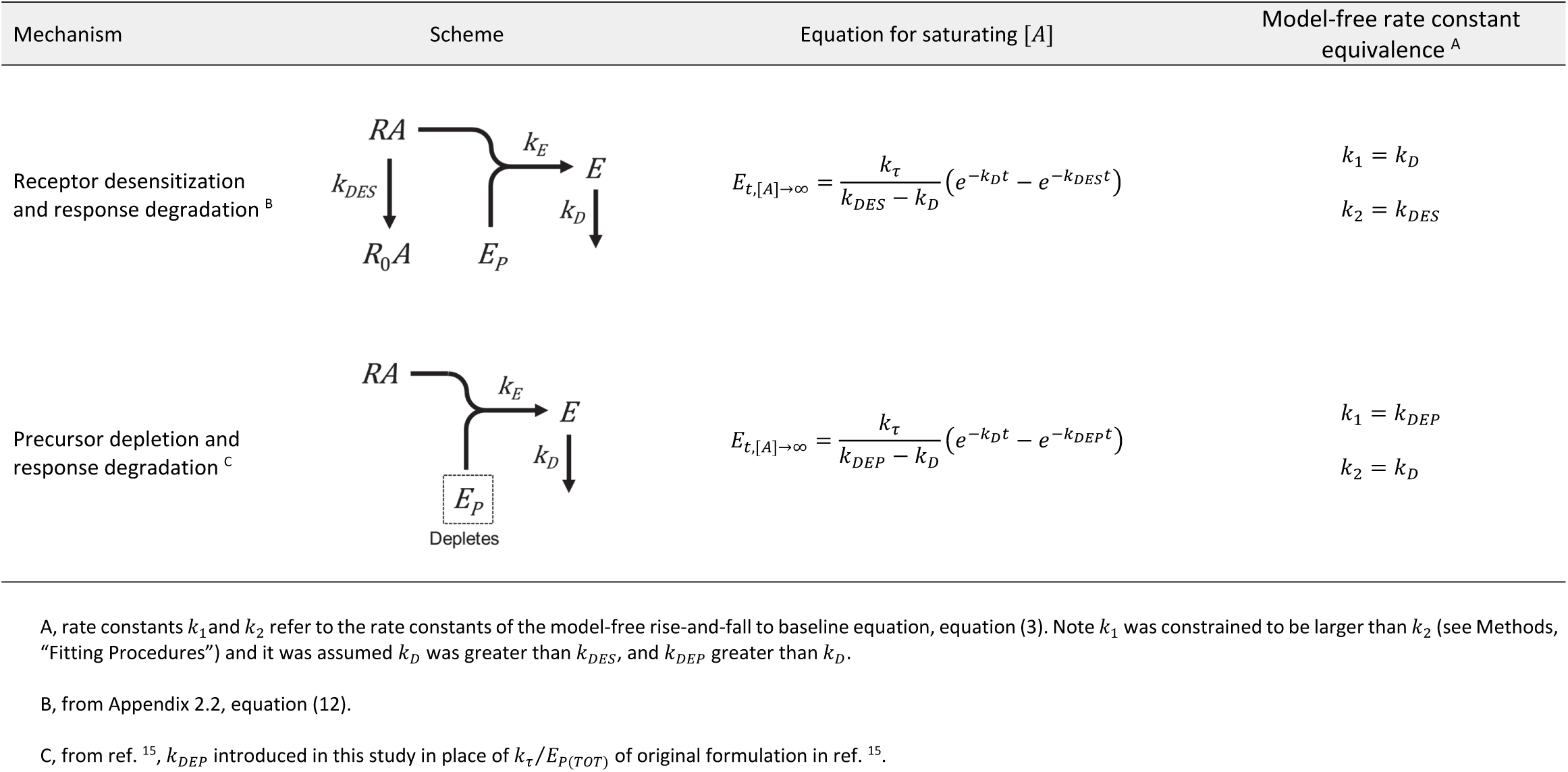
Summary of mechanisms for rise-and-fall to baseline time course profiles.

| Mechanism | Scheme | Equation for saturating $[A]$ | Model-free rate constant equivalence <sup>A</sup> |
| --- | --- | --- | --- |
| Receptor desensitization and response degradation <sup>B</sup> | | $E_{t,[A] \rightarrow \infty} = \frac{k_{\tau}}{k_{DES} - k_D} (e^{-k_D t} - e^{-k_{DES} t})$ | $k_1 = k_D$<br>$k_2 = k_{DES}$ |
| Precursor depletion and response degradation <sup>C</sup> | | $E_{t,[A] \rightarrow \infty} = \frac{k_{\tau}}{k_{DEP} - k_D} (e^{-k_D t} - e^{-k_{DEP} t})$ | $k_1 = k_{DEP}$<br>$k_2 = k_D$ |

A, rate constants $k_1$ and $k_2$ refer to the rate constants of the model-free rise-and-fall to baseline equation, equation (3). Note $k_1$ was constrained to be larger than $k_2$ (see Methods, “Fitting Procedures”) and it was assumed $k_D$ was greater than $k_{DES}$ , and $k_{DEP}$ greater than $k_D$ .
B, from Appendix 2.2, equation (12).
C, from ref. <sup>15</sup>, $k_{DEP}$ introduced in this study in place of $k_{\tau}/E_{P(TOT)}$ of original formulation in ref. <sup>15</sup>.

**Supplemental Table S4.** Summary of mechanisms for rise-and-fall to steady-state time course profiles.

| Mechanism | Scheme | Equation for saturating $[A]$ | Model-free rate constant equivalence <sup>A</sup> |
| --- | --- | --- | --- |
| Receptor desensitization and resensitization with response degradation <sup>B</sup> | | $E_{t,[A] \rightarrow \infty} = \frac{k_{\tau} k_{RES}}{k_D k_{DR}} \times \left[ 1 - \frac{k_{DR}}{k_{DR} - k_D} e^{-k_D t} + \frac{k_D}{k_{DR} - k_D} e^{-k_{DR} t} \right] + \frac{k_{\tau}}{k_{DR} - k_D} (e^{-k_D t} - e^{-k_{DR} t})$ | $k_1 = k_D$<br>$k_2 = k_{DES} + k_{RES}$ |
| Receptor desensitization, desensitized receptor signals, with response degradation <sup>C</sup> | | $E_{t,[A] \rightarrow \infty} = \frac{k_{\tau 2}}{k_D} (1 - e^{-k_D t}) + \frac{k_{\tau 1} - k_{\tau 2}}{k_{DES} - k_D} (e^{-k_D t} - e^{-k_{DES} t})$ | $k_1 = k_D$<br>$k_2 = k_{DES}$ |
| Response degradation to steady-state with precursor depletion <sup>D</sup> | | $E_{t,[A] \rightarrow \infty} = \frac{k_{\tau} k_R}{k_{D(obs)} k_{DEP}} \times \left[ 1 - \frac{k_{DEP}}{k_{DEP} - k_{D(obs)}} e^{-k_{D(obs)} t} + \frac{k_{D(obs)}}{k_{DEP} - k_{D(obs)}} e^{-k_{DEP} t} \right] + \frac{k_{\tau}}{k_{DEP} - k_{D(obs)}} (e^{-k_{D(obs)} t} - e^{-k_{DEP} t})$ | $k_1 = k_{DEP}$<br>$k_2 = k_R + k_D$ |

A, rate constants $k_1$ and $k_2$ refer to the rate constants of the model-free rise-and-fall to steady-state equation, equation (4). Note $k_1$ was constrained to be larger than $k_2$ (see Methods, “Fitting Procedures”) and it was assumed $k_D$ was greater than $k_{DES} + k_{RES}$ ; $k_D$ was greater than $k_{DES}$ ; and $k_{DEP}$ greater than $k_R + k_D$ .
B, from Appendix 2.3.1, equation (13). Note $k_{DR} = k_{DES} + k_{RES}$ .
C, from Appendix 2.3.2, equation (14).
D, from Appendix 2.3.3, equation (16). Note $k_{D(obs)} = k_R + k_D$ .

**Supplementary Table S5.**
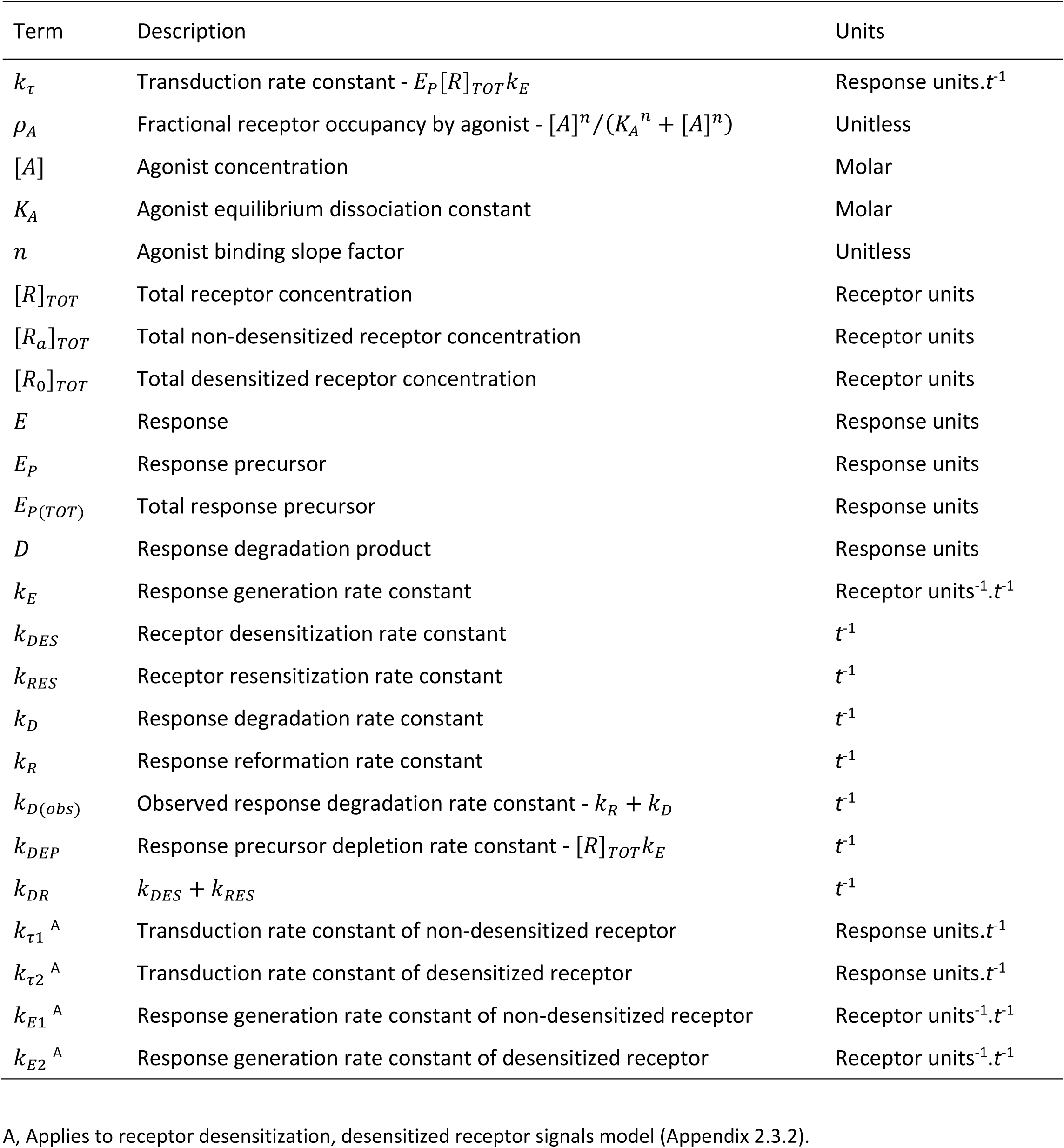
Definition of model parameters

### Model equations list

These equations are loaded into Prism 8 as user-defined equations in a template called “Signaling kinetic mechanism equations” available in the supporting files.

Note in all cases,

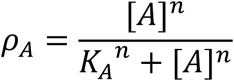

#### No regulation

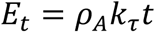

Equation (1) of ref. ^15^

#### No regulation, max A

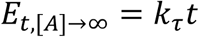

#### Receptor desensitization, max A

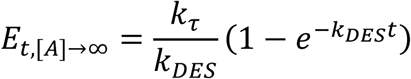

Equation (11) from Appendix 2.1. Note model was formulated only for saturating *[A]*.

#### Response degradation

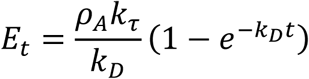

Equation (2) of ref. ^15^

Response degradation, max A

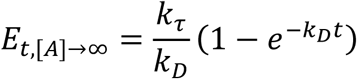

#### Response recycling

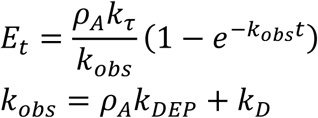

Equation (3) or ref. ^15^, modified so that *k*_*DEP*_ replaces *k*_*r*_*/E*_*P(TOT)*_.

#### Response recycling, max A

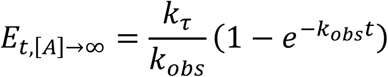

Note *k*_*obs*_ *= k*_*DEP*_ *+ k*_*D*_ but *k*_*DEP*_ and *k*_*D*_ cannot be independently identified with saturating *[A]*.

#### Receptor desensitization & response degradation, max A

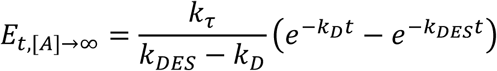

Equation (12) from Appendix 2.2. Note model was formulated only for saturating *[A]*.

#### Precursor depletion & response degradation

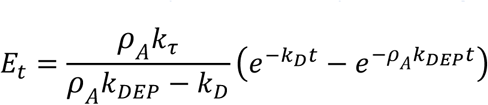

Equation (4) or ref. ^15^, modified so that *k*_*DEP*_ replaces *k*_*r*_*/E*_*P(TOT)*_.

#### Precursor depletion & response degradation, max A

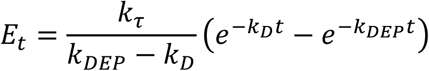

Note, *k*_*DEP*_ assumed to be greater than *k*_*D*_, constrained to be so in fit.

#### Receptor desensitization & resensitization, & response degradation, max A

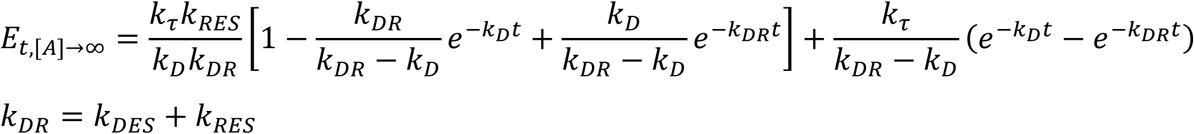

Equation (13) from Appendix 2.3.1. Note model was formulated only for saturating *[A]*.

#### Receptor desensitization, desensitized receptor signals, & response degradation, max A

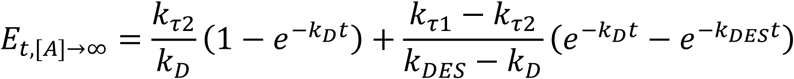

Equation (14) from Appendix 2.3.2. Note model was formulated only for saturating *[A]*.

#### Precursor depletion & response degradation to steady-state

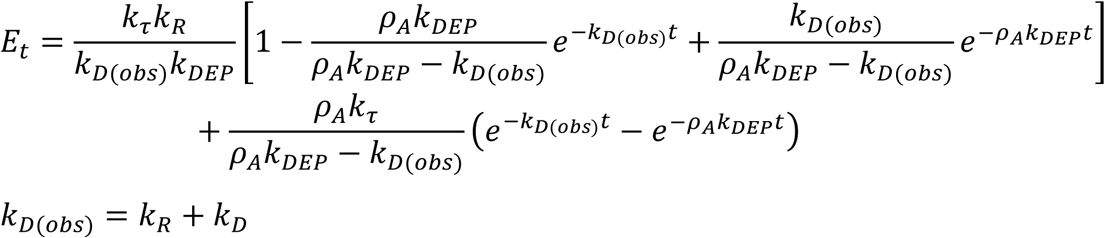

Equation (15) from Appendix 2.3.3.

#### Precursor depletion & response degradation to steady-state, max A

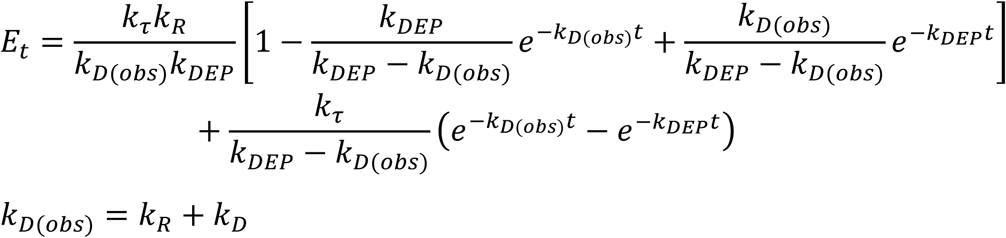

Equation (16) from Appendix 2.3.3.

## Notes

### Competing Interest Statement

The authors have declared no competing interest.

### Summary of Updates

Updated with grant support information.

